# Psychometric validation and clinical correlates of an experiential foraging task

**DOI:** 10.1101/2023.12.28.573439

**Authors:** Aaron N. McInnes, Christi R. P. Sullivan, Angus W. MacDonald, Alik S. Widge

**Affiliations:** Department of Psychiatry & Behavioral Sciences, University of Minnesota, Minneapolis, MN, USA; Department of Psychology, University of Minnesota, Minneapolis, MN, USA

**Keywords:** decision making, reliability, foraging, computational psychiatry, WebSurf

## Abstract

Measuring the function of decision-making systems is a central goal of computational psychiatry. Individual measures of decisional function could be used to describe neurocognitive profiles that underpin psychopathology and offer insights into deficits that are shared across traditional diagnostic classes. However, there are few demonstrably reliable and mechanistically relevant metrics of decision making that can accurately capture the complex overlapping domains of cognition whilst also quantifying the heterogeneity of function between individuals. The WebSurf task is a reverse-translational human experiential foraging paradigm which indexes naturalistic and clinically relevant decision-making. To determine its potential clinical utility, we examined the psychometric properties and clinical correlates of behavioural parameters extracted from WebSurf in an initial exploratory experiment and a pre-registered validation experiment. Behaviour was stable over repeated administrations of the task, as were individual differences. The ability to measure decision making consistently supports the potential utility of the task in predicting an individual’s propensity for response to psychiatric treatment, in evaluating clinical change during treatment, and in defining neurocognitive profiles that relate to psychopathology. Specific aspects of WebSurf behaviour also correlate with anhedonic and externalising symptoms. Importantly, these behavioural parameters may measure dimensions of psychological variance that are not captured by traditional rating scales. WebSurf and related paradigms might therefore be useful platforms for computational approaches to precision psychiatry.

## 1.0 Psychometric validation and clinical correlates of an experiential foraging task

Foraging is a core behaviour in every motile species, requiring an integration of multiple cognitive functions. It requires an organism to accurately monitor the environment and its contingencies, including the availability of resources and the costs and risks of collecting them, whilst assessing one’s homeostatic state (Lima & Bednekoff, 1999; Mobbs et al., 2018; Stephens, 2008; Stephens & Krebs, 1986). Importantly, given that these behaviours have been subject to strong selective pressure over evolution (Stephens, 2008; Stephens & Krebs, 1986), foraging paradigms offer a naturalistic way to understand the decision-making processes that are pertinent to approach problems that the brain evolved to solve (Mobbs et al., 2018). Thus, foraging paradigms can be useful as a lens to study psychopathology, because many mental disorders are thought to fundamentally arise from problems in decision-making systems (Adams et al., 2016; Kishida et al., 2010; Redish, 2013). Neuroeconomic tasks model behavioural strategies that animals adopt and are increasingly being used in psychiatric research as a means to elucidate the relationship between decision-making and psychopathology (Hasler, 2012; Redish et al., 2021).

Previous efforts to translate animal foraging paradigms to human research have often applied secondary reinforcements such as points or money (Kolling et al., 2012; Reynolds & Schiffbauer, 2004; Shenhav et al., 2014). This may impact the validity of these tasks because secondary reward, as opposed to primary consummatory reward, may evoke activity in different neurobiological reward circuitry (Abram et al., 2016). This is pertinent to the validity of these tasks in understanding psychopathology because aberrant reward processing is prevalent transdiagnostically (Zald & Treadway, 2017). The WebSurf task (Abram et al., 2016; and its rodent counterpart, Restaurant Row; Steiner & Redish, 2014; Sweis et al., 2018) is a serial foraging paradigm developed to address this issue. In these tasks, subjects make decisions whether to accept or reject time delays for primary rewards (entertaining videos in WebSurf, flavoured food pellets in Restaurant Row (Abram et al., 2016; Abram, et al., 2019; Abram, et al., 2019; Huynh et al., 2021; Steiner & Redish, 2014; Sweis et al., 2018). In the WebSurf task, participants move sequentially between four video ‘galleries’ and decide whether to invest time from a limited budget in order to receive reward, or forgo the time delay and move to the next video gallery. Importantly, multiple decisional constructs can be derived from task behaviour and behavioural patterns are consistent across human and rodent species (Abram, et al., 2019; Kazinka et al., 2021; Sweis et al., 2018).

Applying foraging paradigms may be useful in establishing neurocognitive profiles that support psychopathology and which are mechanistically relevant, rather than relying solely on clinical rating scales. The establishment of such neurocognitive profiles may facilitate the development of animal models in psychiatry by allowing parallel investigation across species, and offer insights to psychiatric comorbidities by defining deficits that are shared across traditional psychiatric classification (Adams et al., 2016; Huys et al., 2016; Redish et al., 2021; Redish & Gordon, 2016; Robbins et al., 2012). However, the absence of psychometric evidence such as good retest reliability and minimal ceiling and floor effects impedes further progress because one cannot assume that psychophysical tasks accurately capture individual differences in behaviour (Poldrack & Yarkoni, 2016). For example, tasks that are traditionally used to quantify decision-making are typically optimised to index individual domains of cognition, at the expense of capturing the interactions between multiple overlapping constructs. Furthermore, these tasks often demonstrate only modest retest reliability, and show poor inter-individual variability, because they increase the observable differences between experimental conditions and suppress variability between individuals (Dang et al., 2020; Enkavi et al., 2019; Goschke, 2014; Poldrack & Yarkoni, 2016; Redish et al., 2021; Strauss et al., 2006). More naturalistic behavioural tasks, which do not rely on a specific A/B contrast, may be more useful in examining decisional systems which reflect psychopathology (Dang et al., 2020; Enkavi et al., 2019; Poldrack & Yarkoni, 2016), however this hypothesis awaits confirmation.

The WebSurf task holds value as a clinically meaningful tool to examine aberrant decisional processing because it addresses the issues faced by many other neurocognitive tools. However, it has yet to be determined whether behaviour in WebSurf is reliable over time and whether it holds predictive utility for psychopathological symptoms. For rodents, behaviour in this paradigm remains consistent over multiple days of testing (Steiner & Redish, 2014; Sweis et al., 2018) but differs from animal to animal. While humans show between-individual variability consistent with that of animals (Abram, Hanke, et al., 2019; Huynh et al., 2021; Kazinka et al., 2021; Redish et al., 2022; Sweis et al., 2018), within-individual behavioural consistency across multiple days of testing has not been confirmed. Establishing reliability of a measure can indicate both its signal-to-noise ratio and its ability to capture trait constructs. This is important to establish for a task to be used as a valid metric of clinical change (such as that during treatment), as a predictor of response to treatment, and to determine who should receive treatment. Furthermore, variants of the WebSurf task have identified behavioural patterns that are associated with trait externalising features, which is a risk factor for addiction (Abram, Redish, et al., 2019). However, it remains unclear whether task behaviour can predict a wider range of psychopathological symptoms. Finally, for the task to be clinically scalable, it should be valid in an unsupervised environment (Lavigne et al., 2022). Given the primary risk in WebSurf is boredom, unsupervised administration of the task may threaten its computational validity because participants can have unlimited access to distractions that relieve task-evoked boredom (Weydmann et al., 2022). For example, when administered online, some participants demonstrate uneconomical WebSurf task behaviour (preferring longer delays) that may indicate self-distraction (Huynh et al., 2021; Kazinka et al., 2021). Therefore, if the task is to be used as a scalable, clinical tool, it must be validated to determine whether behavioural quantification is repeatable and reliable when participants are unsupervised.

Here, we examined the clinical utility of the WebSurf task in this unsupervised context. Specifically, we tested the stability of task parameters over time, and applied behavioural checks to determine the frequency at which obvious task inattentiveness occurred in an uncontrolled and unsupervised online environment. Finally, we administered a battery of questionnaires assessing symptoms of internalising and externalising psychopathology to determine how patterns of task behaviour may relate to certain psychopathological profiles.

We conducted these in an initial exploratory experiment and validated those findings with a pre-registered confirmatory experiment.

## 2.0 Methods

### 2.1 Participants

Participants (n = 201) were recruited from the online platform Prolific for participation in Experiment One (exploratory sample). To receive an invitation to participate in the study, participants were required to be a resident of the United States and be older than 18 years of age. Participants were also required to have access to a desktop computer or laptop with audio capabilities. Half of these participants completed a single session of the WebSurf task (Group A) and the other half completed repeated sessions (Group B). This study was approved for human subjects research by the University of Minnesota local Institutional

Review Board (STUDY00007274). All participants provided informed consent in accordance with the Declaration of Helsinki.

We recruited a second sample of 200 participants from Prolific to perform validation analyses of our findings. The demographics of the exploratory (Experiment One) and validation (Experiment Two) samples are reported in Table 1.

**Table 1:** Demographics of recruited samples.

|  |  | **Experiment One** |  |  | **Experiment Two** |  |  |
| --- | --- | --- | --- | --- | --- | --- | --- |
|  |  | **Group A**<br>**(n = 101)** | **Group B**<br>**(n = 100)** | **All**<br>**(n = 201)** | **Group A**<br>**(n = 100)** | **Group B**<br>**(n = 100)** | **All**<br>**(n = 200)** |
| **Age** | **Min**<br>**(years)** | 18 | 19 | 18 | 20 | 21 | 20 |
|  | **Max**<br>**(years)** | 65 | 71 | 71 | 78 | 72 | 78 |
|  | **Did not**<br>**specify**<br>**(n)** | 3 | 0 | 3 | 0 | 5 | 5 |
|  | **Mean**<br>**(SD)**<br>**[years]** | 35.06<br>(11.60) | 34.83<br>(12.35) | 34.94<br>(12.48) | 36.47<br>(12.97) | 36.87<br>(11.11) | 36.67<br>(12.07) |
| **Gender**<br>**Identity** | **Female**<br>**(n)** | 49 | 56 | 105 | 41 | 45 | 86 |
|  | **Male (n)** | 49 | 42 | 91 | 58 | 53 | 111 |
|  | **Another**<br>**gender**<br>**(n)** | 2 | 2 | 4 | 0 | 0 | 0 |
|  | **Did not**<br>**specify**<br>**(n)** | 1 | 0 | 1 | 1 | 2 | 3 |
| **Ethnicity** | **% Asian** | 4.95 | 8 | 6.46 | 5 | 9 | 7 |
|  | **% Black** | 12.87 | 7 | 9.95 | 13 | 14 | 13.5 |
|  | **%**<br>**Mixed** | 4.95 | 10 | 7.46 | 8 | 6 | 7 |
|  | **%**<br>**Another** | 2.97 | 6 | 4.48 | 5 | 4 | 4.5 |
|  | **ethnicity** |  |  |  |  |  |  |
|  | **% White** | 67.33 | 68 | 67.66 | 68 | 63 | 65.5 |
|  | **% Did not specify** | 6.93 | 1 | 3.98 | 1 | 4 | 2.5 |

### 2.2 Procedure

The same data collection procedures were conducted for both experiments. After consenting to participate in the study, participants were directed to a battery of self-report clinical rating scales, measuring facets of compulsivity, psychosis, depression, and other related psychopathology. Participants were compensated with $5 USD for completing the questionnaire battery. Twenty-four hours after completing the questionnaire battery, participants were invited to complete a session of the informational foraging task WebSurf. The task had a duration of 30 minutes, and participants received $6 USD for completion of the task. In addition, a random subset of 100 participants (referred to as Group B) were invited to participate in two further sessions of the WebSurf task. This Group was intended to assess reliability of task behaviour. An invitation was sent 24 hours after completion of the first WebSurf session, and another 72 hours after completion of the second WebSurf session. Participants were compensated with $6 and $7 USD for completion of the second and third WebSurf sessions, respectively.

#### 2.2.1 Measures

##### 2.2.1.1 WebSurf Task

The WebSurf task simulates foraging behaviour in humans, in which participants move between four galleries (kittens, dance, bike accidents, landscapes) of videos which function as a reward for investing time (Abram et al. 2016). The task sequence is shown in Figure 1. Upon arrival to a video gallery, participants are informed of a time delay that they must wait for the current gallery’s video to download. This wait duration is randomised from a uniform distribution of 3 – 30 seconds. Participants can choose to accept or reject the offer. If the participant decides to skip, they move to the next video gallery, where they are given another offer which they must choose to accept or reject. If the participant decides to accept the offer, they must wait for the reward. During the wait-period, participants can choose to quit out of the wait and move to the next gallery. If participants successfully wait for the entire duration, a video from the current gallery plays for approximately four seconds. At the completion of each video, participants are asked to rate the video using a five-star rating system, where one star = extremely dislike and five stars = extremely like. Once the video is rated, participants move to the next video gallery and are presented with another offer. The four video galleries presented in the task were consistent across sessions, but unique videos within each gallery were presented in each session.

**Figure 1.**
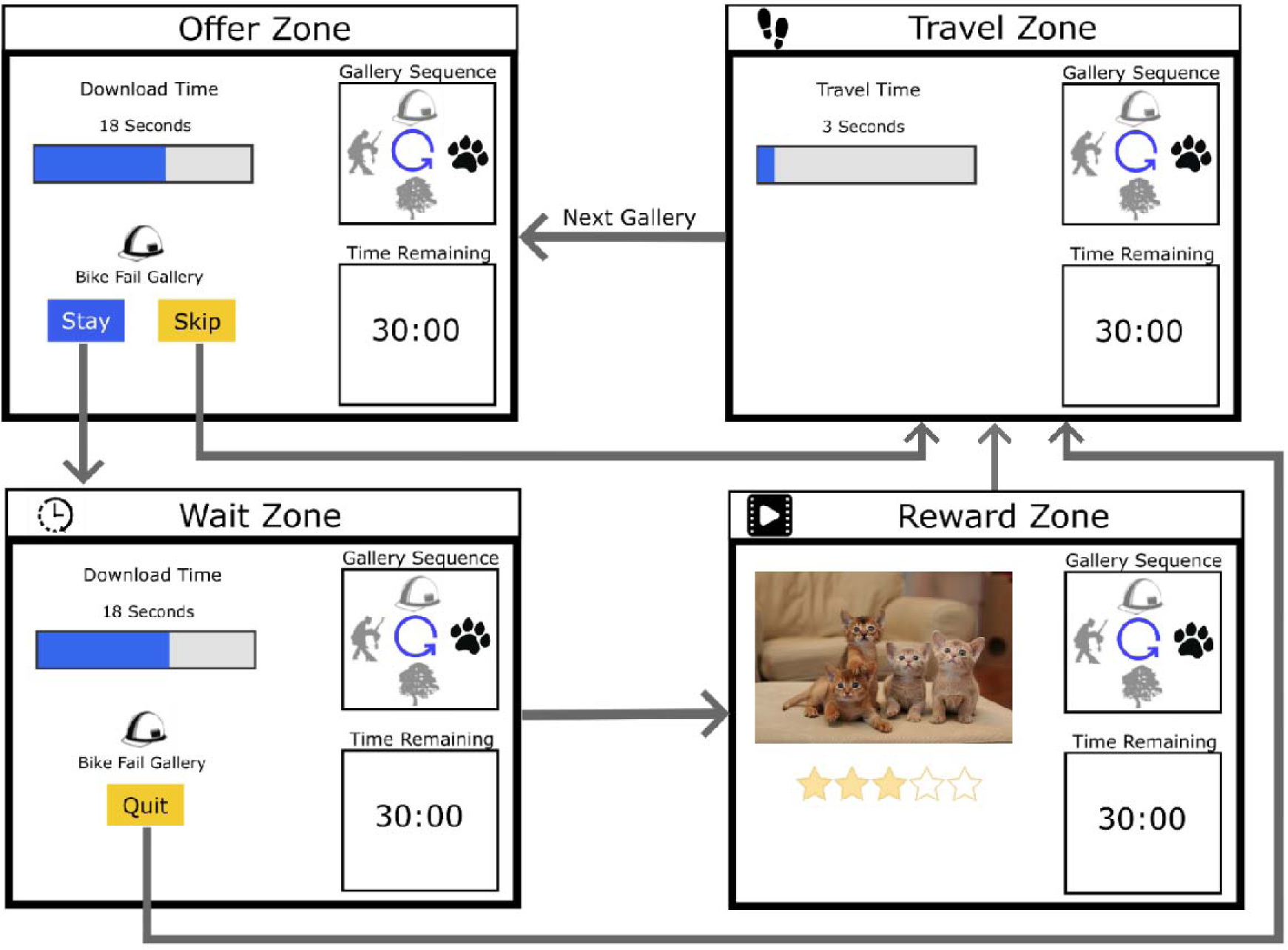
Sequence of the WebSurf task. Participants start in the Offer Zone, where they are presented the time delay they must wait for reward. If they accept the offer, the move to the Wait Zone. If they reject the offer, they move to the Travel Zone. In the Wait Zone, participants must wait the accepted duration of time (3 – 30 seconds) before they move to the Reward Zone. If participants decide to quit, they move to the Travel Zone. Once the duration has ended in the Wait Zone, participants are presented a video from the current video gallery.

At the completion of the video, participants provide a rating from 1 – 5 stars and move to the Travel Zone. In the Travel Zone, participants must wait three seconds before moving to the next video gallery.

The core measures of behaviour we extracted from the task are decision threshold, probability of accepting offers (P[Accept]), average video rating, average decision latency, and number of threshold violations. Behavioural parameters such as decision thresholds, P(Accept), and video ratings are clinically relevant constructs to measure because they assess responses to positive motivational contexts, including reward responsiveness and valuation, which are implicated in a range of psychiatric disorders (Barch et al., 2009). Decision latencies can reflect the engagement of deliberative decisional processes as individuals evaluate options and their outcomes (Pleskac et al., 2019). Reaction times may also indicate the efficiency of decisional systems, and the engagement of impulsive or habitual decision strategies. The number of threshold violations is also a relevant behavioural parameter in assessing the optimality of decisions in maximising the ratio of investment to reward, and relates to impulsive decisional styles. An outline of these parameters is provided in Table 2.

**Table 2.**
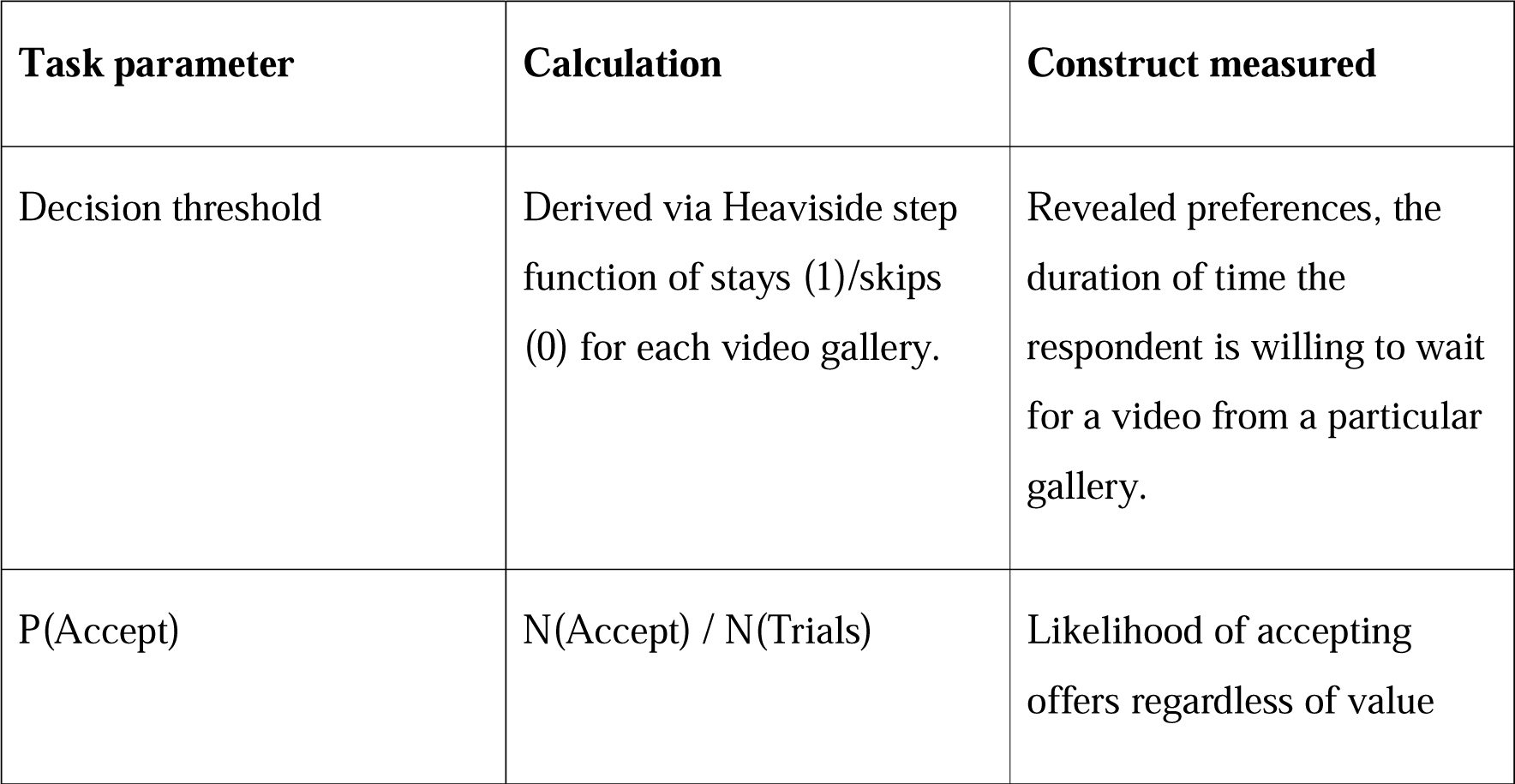
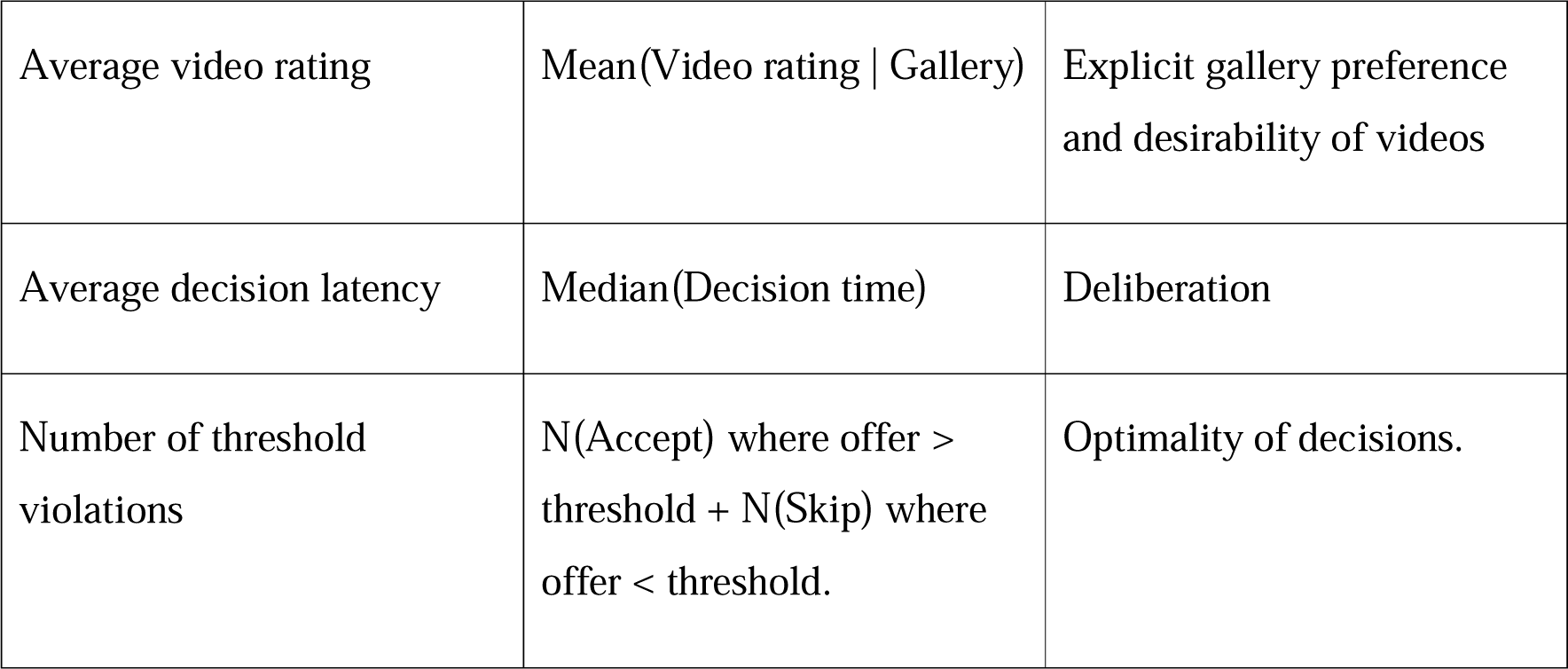
Definition of task parameters.

##### 2.2.1.2 Questionnaire Battery

A battery of previously validated self-report rating scales was administered to participants after initial recruitment into the study. The questionnaire battery included the Altman Self-Rating Mania Scale (ASRM; Altman et al., 1997), the Alcohol Use Disorders Identification Test (AUDIT; Saunders et al., 1993), the Aberrant Salience Inventory (ASI; Cicero et al., 2010), the Centre for Epidemiological Studies – Depression (CES-D; Radloff, 2016), the Daily Sessions, Frequency, Age of Onset, and Frequency of Cannabis Use (DFAQ-CU; Cuttler & Spradlin, 2017), the Eating Atttitudes Test (EAT-26; Garner et al., 1982), the Mini Mood and Anxiety Symptoms Questionnaire (Mini-MASQ; Watson et al., 1995), the Obsessive Compulsive Inventory – Revised (OCI-R; Foa et al., 2002), the Snaith-Hamilton Pleasure Scale (SHAPS; Snaith et al., 1995), the Barratt Impulsivity Scale (BIS-11; Patton et al., 1995), the Schizotypal Personality Questionnaire – Brief (SPQ-B; Raine & Benishay, 1995), the Temporal Experience of Pleasure Scale (TEPS; Gard et al., 2006), and the World Health Organization Alcohol, Smoking, and Substance Involvement Screening Test (WHO-ASSIST; WHO ASSIST Working Group, 2002). These data were used to assess the clinically relevant predictive utility of task behavioural parameters. To this end, we examined whether task behaviour is predictive of symptomology, as measured by individual questionnaire scores, and of psychopathological profiles, as measured by a principal component analysis (PCA) of questionnaire scores. The questionnaires were administered in a fixed order for all participants, with attention-check items interspersed within questionnaire items. Participants were excluded from the study if they failed more than one attention check item in the questionnaire battery. Subject responses to a particular scale were discarded if its corresponding attention-check item was failed. Descriptions of each measure and their subscales are outlined in Supplementary Table S1.

Questionnaire items were reverse scored where necessary and scored items were summed to produce a total score for each questionnaire. The exception to this was the DFAQ-CU, for which the three subscales measure cannabis use on three different scales of measurement (frequency, age of onset, and quantity). As such, items were transformed by computing z-scores within each subscale and summed to produce a total score.

#### 2.2.2 Statistical Analyses

All data processing and analyses were conducted using R (R Core Team, 2016). Prior to analysing behavioural data from the WebSurf task, we screened data for evidence that subjects were inattentive during their participation (see Supplementary Materials). Once total scores were calculated for each questionnaire and WebSurf task parameters were derived, we subjected these data to a series of exploratory analyses in Experiment One. First, we examined the reliability and stability of behaviour in the WebSurf task over repeated administrations. These repeatability tests were conducted with the subset of participants who completed three sessions of the task (Group B). To evaluate reliability of task behaviour, we calculated intraclass correlations (ICCs) of WebSurf task parameters with two-way mixed effects and based on absolute agreement (ICC(2,1); Koo & Li, 2016; McGraw & Wong, 1996; Shrout & Fleiss, 1979). We calculated ICCs for all session combinations (i.e. across all sessions, across the first and second session, across the first and third session, and across the second and third session). Based on criteria proposed by Koo and Li (2016), reliability was deemed adequate if the ICC of a parameter was > 0.75.

In addition, we examined whether there is a systematic change in task parameters with repeated task administration, which may indicate that participants satiate to the task. If task parameters are stable over repeated administrations, then the main effect of session number should not be significant. Given that this proposition relies on confirmation of the null hypothesis, we evaluated it with a series of Bayesian linear mixed-effects models, with WebSurf task parameters (decision thresholds, P(Accept), average video ratings, median decision latencies, and number of threshold violations) as response variables, with session number and category rank as predictor variables, and subject IDs as a random effect. We calculated BF_01_ values to evaluate evidence for the null over the alternative hypothesis and interpreted these based on the criteria outlined by Jeffreys (2006).

For the subsequent analyses, we included all first-session data across Groups A and B. We evaluated hedonic reliability within sessions by calculating correlations between different indices of gallery preference (gallery decision thresholds, average video ratings for each gallery, and explicit gallery rankings). For each combination of these preference indices, we calculated Kendall’s T*_b_* and determined that hedonic reliability was adequate if these values were > 0.3 (Schaeffer & Levitt, 1958).

To determine profiles of psychiatric symptoms which cut across traditional psychiatric classification, we subjected our questionnaire scores to principal component analysis (PCA). Total scores from each questionnaire were normalized via z-transformation prior to running PCA. The most important PCs were selected from the PCA by evaluating the eigenvalues and percentage of questionnaire score variance accounted for by each PC. The PCA rotations were used to calculate individual subject scores for each of the PCs we extracted from our questionnaire data. The resulting components extracted from the PCA are presented in the Supplementary Materials. These PC scores were then used as predictors of WebSurf task behaviour in following analyses. We also ran a series of least absolute shrinkage and selection operator (LASSO) models to identify specific symptom patterns (questionnaire scores) which were significant predictors of task behaviour.

We ran a series of tests to evaluate how psychiatric symptomology can predict behaviour in the WebSurf task. To assess how study drop out may have impacted the representativeness of these data, we evaluated the probability of completing all parts of the study as a function of psychiatric symptomology by running logistic regressions with study completion as the dependent variable. Predictive questionnaire scores were identified via LASSO and predictive PCs were identified by examining the significance of their main effects on study completion. We also evaluated predictors of behavioural reliability over time by entering the standard deviations of task behavioural parameters across all three sessions of the task as response variables in a series of linear models, with predictive questionnaire scores (as identified by LASSO) entered as predictor variables. PC scores were also entered separately into linear models to evaluate which PCs were significant predictors of study completion.

Given that standard deviations are right-skewed, we normalised them via z-transformation prior to entering them as response variables in the linear models. Predictors of hedonic reliability were also evaluated. After calculating individual subject correlations between each of decision thresholds x average video ratings, decision thresholds x explicit gallery rankings, and average video ratings x explicit gallery rankings, we entered these correlation coefficients as response variables into a series of linear models, with either significant questionnaire predictors or the PC scores as predictor variables. Finally, we directly entered behavioural parameters from the WebSurf task as response variables in a series of linear models and identified predictive questionnaire scores via LASSO and evaluated the main effects of each of the PCs on the response variable. For all frequentist tests, statistical significance was determined at α = 0.05 and post hoc tests were corrected for multiple comparisons via the false discovery rate method (Benjamini & Hochberg, 1995). Linear mixed-effects models were conducted with Satterthwaite’s approximation for degrees of freedom (Kuznetsova et al., 2017).

In Experiment Two, we performed a validation of the effects we observed in Experiment One. As such, we conducted the same analyses as Experiment One, but rather than using LASSO regression for variable selection, we directly tested the effects observed in Experiment One. These analyses were pre-registered and the registration report is available at https://osf.io/8h6tz.

## 3.0 Results

### 3.1 Study drop out

First, we report the number of participants that did not complete all parts of the study. One source of attrition was due to the longitudinal nature of the study, with some participants failing to complete parts of the study within the allocated timeframe. In addition, participants were rejected from the study if they failed our online attention checking procedure (see Supplementary Materials). Tables 3 and 4 outline the number of participants in each group who were invited to complete each part of Experiment One (exploratory sample) and Experiment Two (validation sample) and the number of task failures in each.

**Table 3:** Group A sample size – Experiments One and Two.

|  | **Completed**<br>**questionnaire**<br>**battery** | **Questionnaire**<br>**responses**<br>**rejected due**<br>**to failed**<br>**attention**<br>**checks** | **Invited to**<br>**complete**<br>**WebSurf** | **Returned**<br>**and**<br>**started**<br>**WebSurf** | **Failed**<br>**WebSurf** | **Completed**<br>**WebSurf** |
| --- | --- | --- | --- | --- | --- | --- |
| **N**<br>**(Exp.**<br>**1)** | 101 | 1 | 100 | 94 | 27 | 67 |
| **N**<br>**(Exp.**<br>**2)** | 100 | 0 | 100 | 84 | 29 | 55 |

**Table 4:**
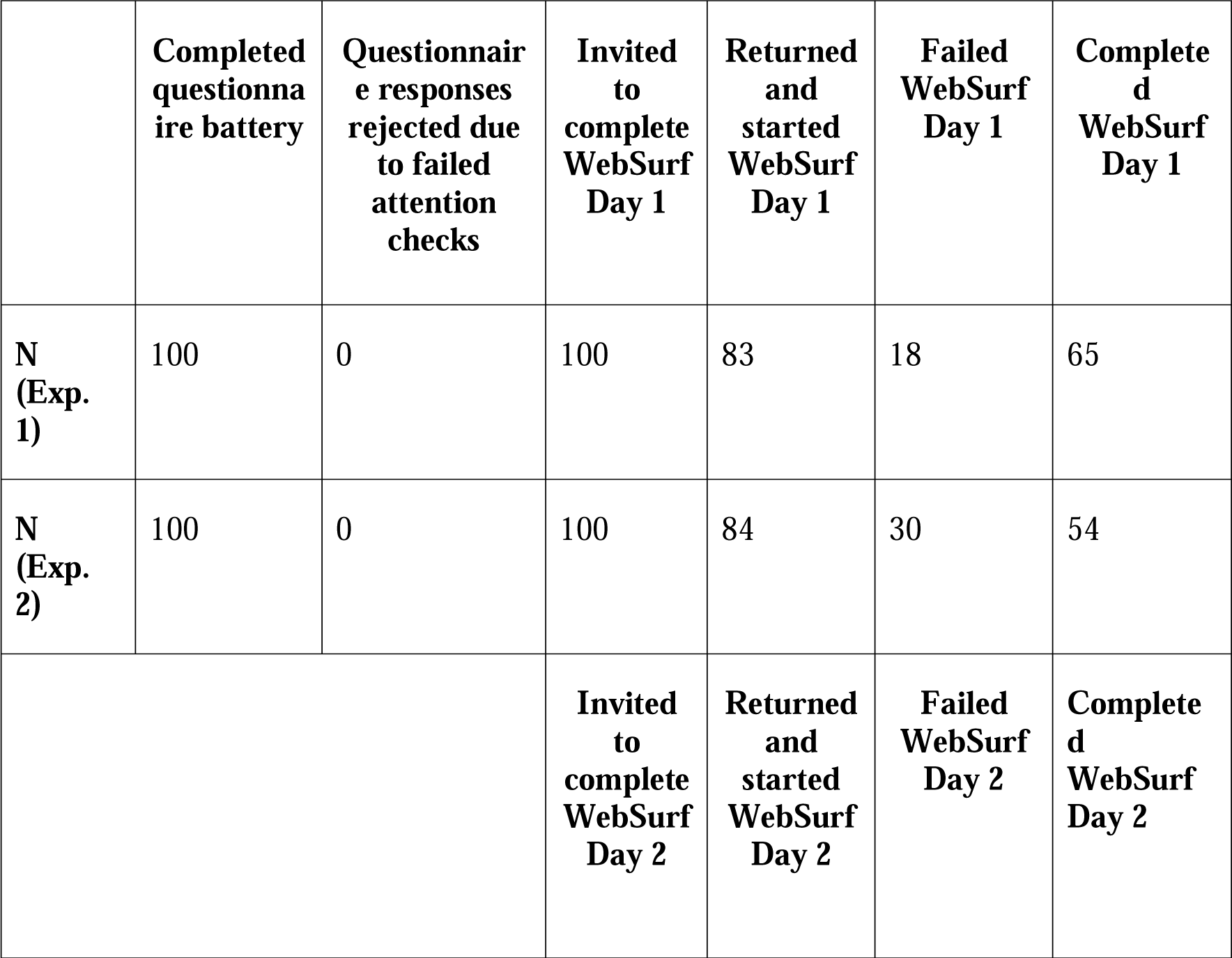
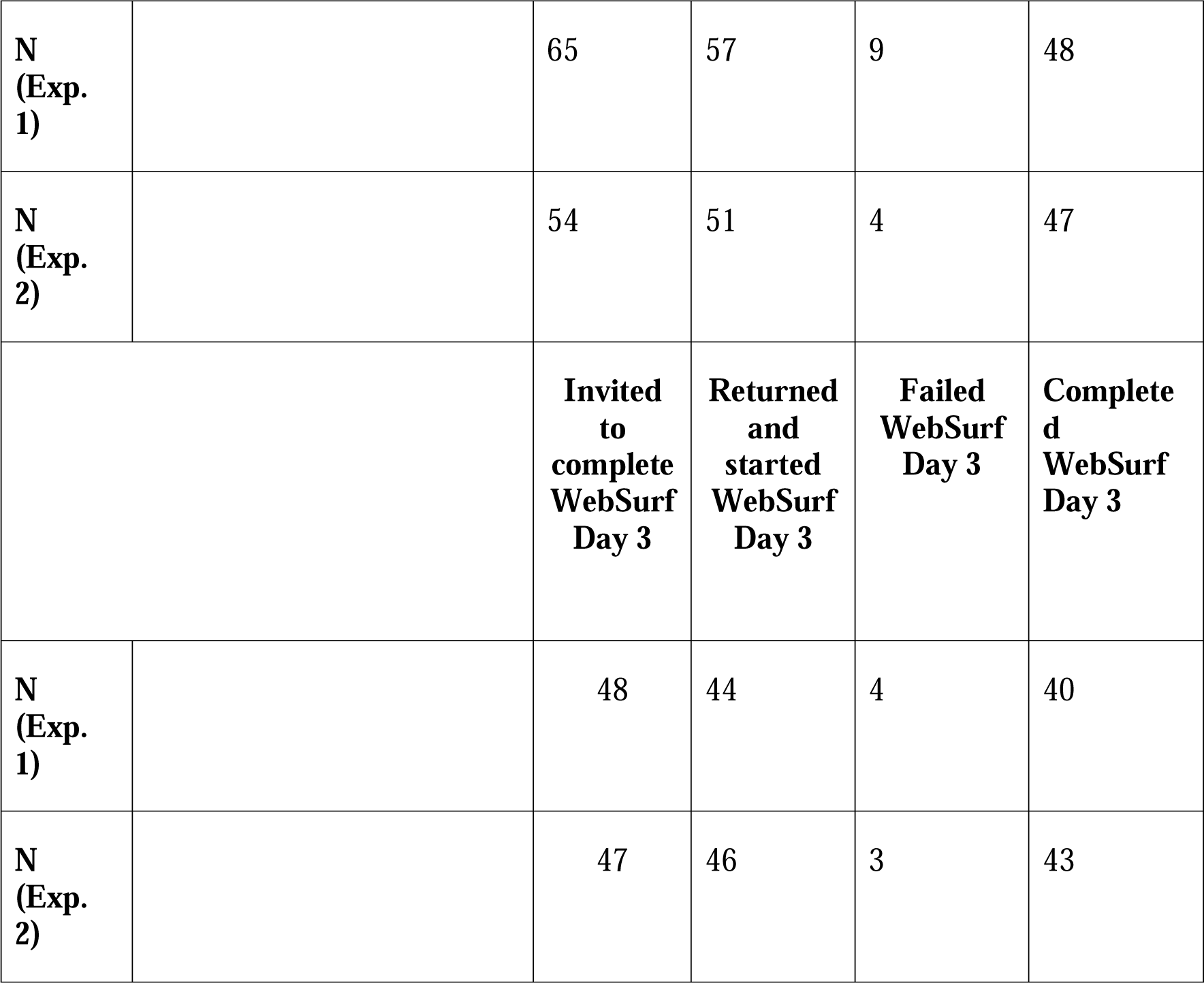
Group B sample size – Experiments One and Two.

|  | **Completed**<br>**questionnaire**<br>**battery** | **Questionnaire**<br>**responses**<br>**rejected due**<br>**to failed**<br>**attention**<br>**checks** | **Invited**<br>**to**<br>**complete**<br>**WebSurf**<br>**Day 1** | **Returned**<br>**and**<br>**started**<br>**WebSurf**<br>**Day 1** | **Failed**<br>**WebSurf**<br>**Day 1** | **Completed**<br>**WebSurf**<br>**Day 1** |
| --- | --- | --- | --- | --- | --- | --- |
| **N**<br>**(Exp.**<br>**1)** | 100 | 0 | 100 | 83 | 18 | 65 |
| **N**<br>**(Exp.**<br>**2)** | 100 | 0 | 100 | 84 | 30 | 54 |
|  |  |  | **Invited**<br>**to**<br>**complete**<br>**WebSurf**<br>**Day 2** | **Returned**<br>**and**<br>**started**<br>**WebSurf**<br>**Day 2** | **Failed**<br>**WebSurf**<br>**Day 2** | **Completed**<br>**WebSurf**<br>**Day 2** |

|  |  |  |  |  |  |
| --- | --- | --- | --- | --- | --- |
| **N**<br>**(Exp.**<br>**1)** |  | 65 | 57 | 9 | 48 |
| **N**<br>**(Exp.**<br>**2)** |  | 54 | 51 | 4 | 47 |
|  |  | **Invited**<br>**to**<br>**complete**<br>**WebSurf**<br>**Day 3** | **Returned**<br>**and**<br>**started**<br>**WebSurf**<br>**Day 3** | **Failed**<br>**WebSurf**<br>**Day 3** | **Complete**<br>**d**<br>**WebSurf**<br>**Day 3** |
| **N**<br>**(Exp.**<br>**1)** |  | 48 | 44 | 4 | 40 |
| **N**<br>**(Exp.**<br>**2)** |  | 47 | 46 | 3 | 43 |

### 3.2 Task Behavioural Parameters

#### 3.2.1 Behavioural Reliability

We examined decision thresholds, P(Accept), average video ratings, median decision times, and number of threshold violations across video gallery ranks and across sessions for the sample of participants who completed all three sessions of WebSurf. Prior to behavioural analysis, we screened data for poor task engagement via our behavioural screening criteria. In total, using these criteria, 11 participants were rejected from Experiment One and eight participants were rejected from Experiment Two.

To examine the reliability of task behavioural parameters over multiple sessions, we calculated ICCs for each combination of sessions. These were calculated for participants that completed all sessions of the task. ICCs of decision thresholds, P(Accept), average video ratings, median decision times, and number of threshold violations are plotted in Figure 2.

**Figure 2.**
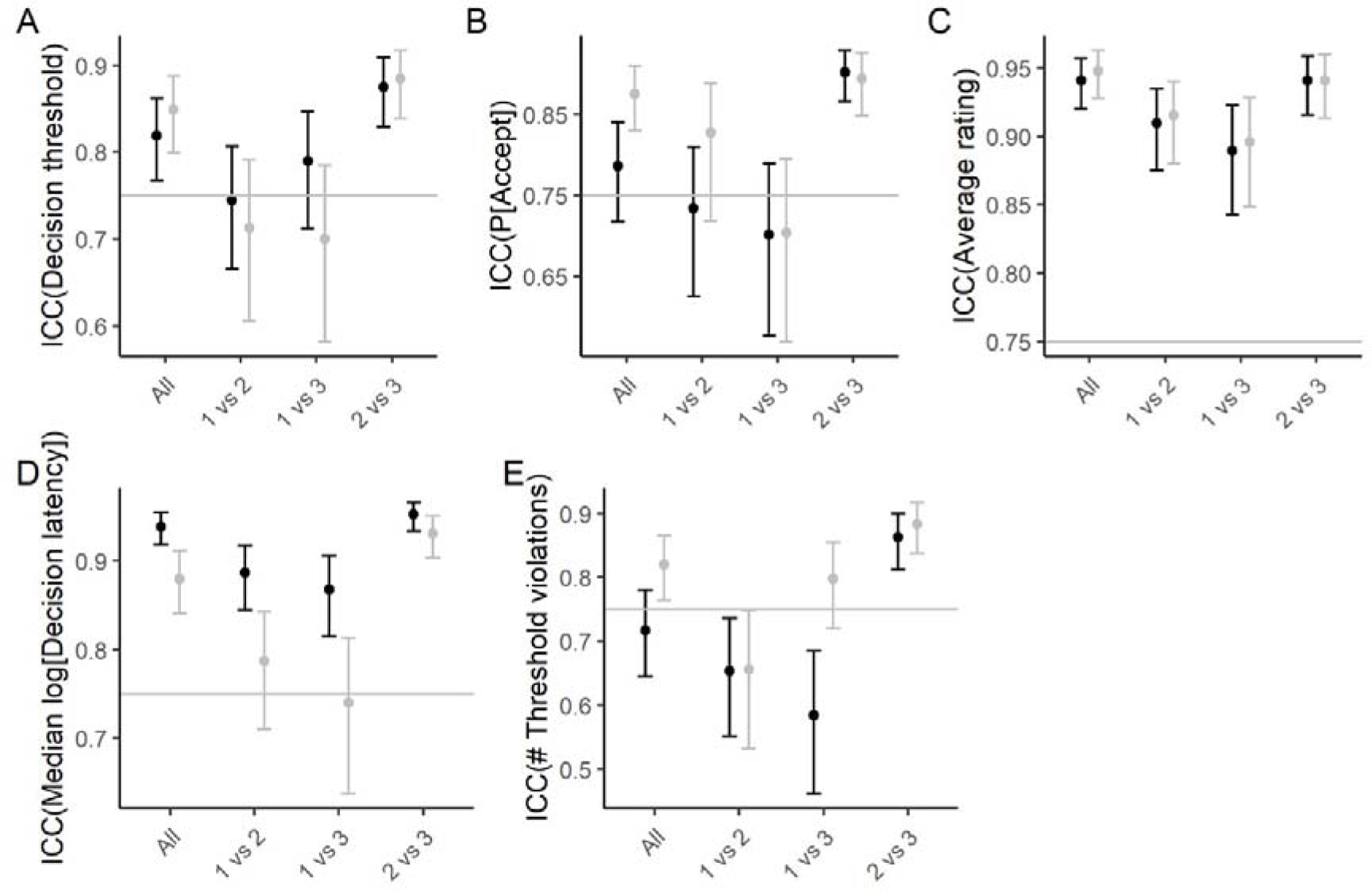
Intraclass correlations (ICCs) of behavioural parameters across sessions of the WebSurf task. Black points represent ICCs from Experiment One. Grey points represent ICCs from Experiment Two. Error bars represent the 95% confidence interval.

ICCs of behavioural parameters between sessions and across video gallery rankings in Experiment One are plotted in Supplementary Figure S3. Supplementary Figure S4 shows these ICCs for Experiment Two. In Experiment One, the most reliable parameters were average video rating and median decision latency, whereas P(Accept) and the number of threshold violations generally did not meet reliability criteria. Also of note is the fact that ICCs were consistently > .75 across the second and third sessions. This suggests that task behaviour becomes more stable after initial exposure to the task. The ICC patterns in Experiment One generally replicated in Experiment Two (see Figure 2). However, decision latencies generally showed poorer reliability in Experiment Two in comparison to Experiment One.

#### 3.2.2 Behaviour over repeated sessions

We further ran a series of linear mixed-effects models to examine whether behavioural parameters drift systematically across repeated administrations of the task and whether there is variance in parameters as a function of gallery preference. Such a systematic change in behaviour over repeated administrations may indicate that participants are prone to satiation after repeated exposure to the task. We report the results of these tests in Table 5.

**Table 5.** Results of linear mixed-effects models of task behavioural parameters as a function of session number and gallery rank, across Experiments One and Two.

| Experiment | Behavioural parameter | Predictor | Degrees of freedom | *F* | *p* |
| --- | --- | --- | --- | --- | --- |
| 1 | Decision threshold | Gallery rank | 3, 401.06 | 79.39 | < .001 |
| 2 |  |  | 3, 403.16 | 113.81 | < .001 |
| 1 |  | Session number | 2, 404.76 | 1.30 | .273 |
| 2 |  |  | 2, 423.13 | 0.94 | .391 |
| 1 |  | Gallery rank<br>x Session<br>number | 6, 401.06 | 0.94 | .466 |
| 2 |  |  | 6, 403.16 | 0.33 | .921 |
| 1 | P(Accept) | Gallery rank | 3, 400.97 | 107.84 | < .001 |
| 2 |  |  | 3, 402.92 | 171.36 | < .001 |
| 1 |  | Session<br>number | 2, 404.71 | 2.85 | .059 |
| 2 |  |  | 2, 423.66 | 2.06 | .130 |
| 1 |  | Gallery rank<br>x Session<br>number | 6, 400.97 | 0.59 | .738 |
| 2 |  |  | 6, 402,92 | 0.21 | .975 |
| 1 | Average<br>video rating | Gallery rank | 3, 383.48 | 136.61 | < .001 |
| 2 |  |  | 3, 383.11 | 190.70 | < .001 |
| 1 |  | Session<br>number | 2, 385.49 | 0.57 | .568 |
| 2 |  |  | 2, 398.41 | 0.56 | .569 |
| 1 |  | Gallery rank<br>x Session<br>number | 6, 385.15 | 0.23 | .968 |
| 2 |  |  | 6, 283.50 | 0.86 | .527 |
| 1 | Median | Gallery rank | 3, 401.11 | 0.90 | .442 |
| 2 | decision<br>latency |  | 3, 403.94 | 8.10 | < .001 |
| 1 |  | Session<br>number | 2, 401.96 | 0.56 | .572 |
| 2 |  |  | 2, 411.16 | 9.99 | < .001 |
| 1 |  | Gallery rank<br>x Session<br>number | 6, 401.11 | 0.31 | .929 |
| 2 |  |  | 6, 403.94 | 0.32 | .931 |
| 1 | Number of<br>threshold<br>violations | Gallery rank | 3, 401 | 10.49 | < .001 |
| 2 |  |  | 3, 403.90 | 15.07 | < .001 |
| 1 |  | Session<br>number | 2, 403.79 | 2.13 | .120 |
| 2 |  |  | 2, 416.39 | 2.0 | .137 |
| 1 |  | Gallery rank<br>x Session<br>number | 6, 401 | 0.77 | .599 |
| 2 |  |  | 6, 403.90 | 1.58 | .152 |

Decision thresholds did not change systematically across repeated administrations of the task, with no significant main effect of session number or interaction of session number with video gallery. A Bayesian linear model supported this conclusion, with very strong evidence against an effect of session number on decision thresholds, BF_01_ = 32.20. There was variation in decision thresholds between galleries, as indicated by the significant main effect of video gallery rank. Decision thresholds decreased with increasing gallery preference, as would be expected by basic economic principles (*M_RANK1_* = 28.02, *SD_RANK1_* = 6.38; *M_RANK2_* = 25.69, *SD_RANK2_* = 5.97; *M_RANK3_* = 23.3, *SD_RANK3_* = 7.1; *M_RANK4_* = 14.88, *SD_RANK4_* = 10.01). Pairwise contrasts indicated a significant difference in decision threshold between all gallery ranks. In Experiment Two, these results replicated. Decision thresholds were stable across sessions, as indicated by no main effect of session number and no interaction of session number with video gallery. Again, there was variance in decision thresholds between galleries, with a significant main effect of gallery rank. Decision thresholds decreased with increasing gallery preference (*M_RANK1_* = 27.83, *SD_RANK1_* = 6.49; *M_RANK2_* = 26.20, *SD_RANK2_* = 7.00; *M_RANK3_* = 21.27, *SD_RANK3_* = 8.63; *M_RANK4_* = 11.13, *SD_RANK4_* = 10.15). Pairwise contrasts indicated a significant difference in decision threshold between all gallery ranks, except for the difference between ranks one and two. Supplementary Figures S1 – S2 show the parameters as a function of session number and gallery rank for Experiments One and Two.

In both Experiment One and Experiment Two, the probability of accepting offers did not vary systematically across sessions. A Bayesian linear model supported the stability of P(Accept) across sessions, providing strong evidence against an effect of session number on P(Accept), BF_01_ = 18.81 (Experiment One), BF_01_ = 28.47 (Experiment Two). The probability of accepting offers did vary across video gallery ranks. In Experiment One the probability of accepting offers increased with increasing gallery preference (*M_RANK1_* = .95, *SD_RANK1_* = .19; *M_RANK2_* = .86, *SD_RANK2_* = .17; *M_RANK3_* = .77, *SD_RANK3_* = .21; *M_RANK4_* = .49, *SD_RANK4_* = .31) and pairwise contrasts indicated a significant difference in the probability of accepting offers between all gallery ranks. This was replicated in Experiment Two (*M_RANK1_* = .90, *SD_RANK1_* = .19; *M_RANK2_* = .83, *SD_RANK2_* = .20; *M_RANK3_* = .66, *SD_RANK3_* = .25; *M_RANK4_* = .31, *SD_RANK4_* = .28).

Average video ratings were also stable across sessions. In Experiment One, this was supported by a Bayesian linear model which indicated strong evidence against change in video ratings as a function of session number, BF_01_ = 17.80. The stability of video ratings across sessions was replicated in Experiment Two, BF_01_ = 32.61. A linear model indicated average video ratings differed across gallery ranks, showing an increase with increasing gallery preference in both Experiment One (*M_RANK1_* = 4.09, *SD_RANK1_* = 0.54; *M_RANK2_* = 3.74, *SD_RANK2_* = 0.54; *M_RANK3_* = 3.17, *SD_RANK3_* = 0.66; *M_RANK4_* = 2.54, *SD_RANK4_*= 0.87), and Experiment Two (*M_RANK1_* = 4.05, *SD_RANK1_* = 0.63; *M_RANK2_* = 3.73, *SD_RANK2_* = 0.61; *M_RANK3_* = 3.01, *SD_RANK3_* = 0.63; *M_RANK4_* = 2.27, *SD_RANK4_* = 0.79). Pairwise contrasts indicated a significant difference in average video ratings between all gallery ranks, and this was replicated in Experiment Two.

In Experiment One, log normalised median decision times showed no main effect of session number and no interaction of session number with gallery preference. The stability of decision times across sessions was supported by a Bayesian linear model which yielded very strong evidence against a change in decision times as a function of session number, BF_01_ = 32.61. The main effect of gallery preference was not significant, indicating decision latencies also did not differ systematically as a function of gallery preference. However, in Experiment Two, there was a systematic change in decision latencies across sessions. There was a significant difference in median decision latencies between session one (*M* = 7.46, *SD* = 0.18) and session three (*M* = 7.37, *SD* = 0.20, *p* < .001), between session two (*M* = 7.43, *SD* = 0.21) and session three (*p* = .003), but not between session one and session two (*p* = .132). In addition, in contrast to Experiment One, there was a main effect of gallery preference in Experiment Two. Decision latencies decreased with gallery preference (*M_RANK1_* = 7.38, *SD_RANK1_* = 0.17; *M_RANK2_*= 7.41, *SD_RANK2_* = 0.19; *M_RANK3_* = 7.42, *SD_RANK3_* = 0.19; *M_RANK4_* = 7.49, *SD_RANK4_* = 0.20). The difference in median decision latency was significant between gallery ranks one and four, ranks two and four, and ranks three and four.

The number of threshold violations appeared stable across sessions in Experiment One. Neither the main effect of session number, nor the interaction of session number with gallery rank, was significant. A Bayesian model indicated stability across sessions, with strong evidence against an effect of session number on threshold violations, BF_01_ = 14.91. The reliability of threshold violations across sessions was replicated in Experiment Two, and the Bayesian model provided substantial evidence against a change in the number of threshold violations as a function of session number, BF_01_ = 4.69. There was variation between galleries , as indicated by a decrease of the number of threshold violations as gallery preference increased in Experiment One (*M_RANK1_* = 1.06, *SD_RANK1_* = 1.96; *M_RANK2_* = 1.81, *SD_RANK2_* = 2.09; *M_RANK3_* = 2.04, *SD_RANK3_*= 2.01; *M_RANK4_* = 2.50, *SD_RANK4_* = 2.47).

Pairwise contrasts indicated a significant difference in threshold violations between all gallery ranks except for the difference between ranks two and three (*p* = .424), and between ranks three and four (*p* = .087). This was replicated in Experiment Two, with threshold violations decreasing as gallery preference increased, (*M_RANK1_* = 1.85, *SD_RANK1_* = 2.35; *M_RANK2_* = 2.12, *SD_RANK2_* = 2.30; *M_RANK3_* = 3.52, *SD_RANK3_* = 2.99; *M_RANK4_* = 3.49, *SD_RANK4_* = 2.91). In Experiment Two the difference in the number of threshold violations was significant between all gallery ranks except for the difference between ranks one and two and between ranks three and four.

### 3.3 Hedonic Reliability

We determined each participant’s ranking of each video gallery based on decision threshold, average video rating, and explicit gallery rankings. With each combination of these variables, we calculated Kendall’s T*_b_* to determine hedonic reliability. There was a significant relationship between average video rating rank and decision threshold rank, τ*_b_* [95% CI] = .48 [.41, .55], *p* < .001, between average video rating rank and explicit gallery rank,τ*_b_* [95% CI] = .69 [.64, .73], *p* < .001, and between decision threshold rank and explicit gallery rank, τ*_b_* [95% CI] = .49 [.43, .56], *p* < .001. The frequency of congruency and correlations between revealed and stated preferences are shown in Figure 3.These results replicated in Experiment Two - average video rating rank and decision threshold rank, τ*_b_* [95% CI] = .49 [.41, .58], *p* < .001, average video rating rank and explicit gallery rank,τ*_b_* [95% CI] = .70 [.63, .76], *p* < .001, and decision threshold rank and explicit gallery rank, τ*_b_* [95% CI] = .51 [.43, .60], *p* < .001, again showed hedonic reliability.

**Figure 3.**
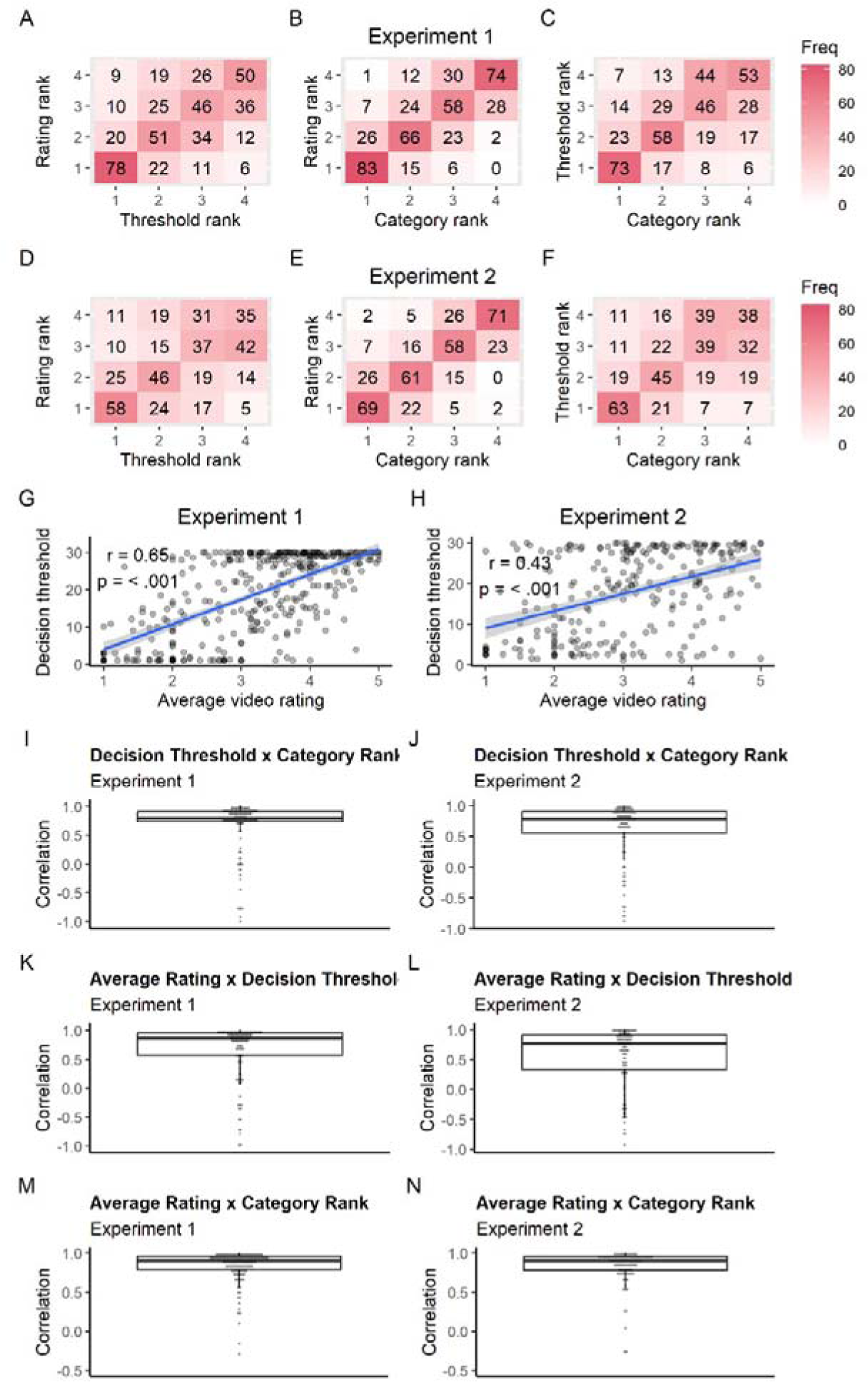
Hedonic reliability of revealed and stated preferences. (A-C) Matrix of revealed and stated preference congruency in Experiment 1. (D-F) Matrix of revealed and stated preference congruency in Experiment 2. (G-H) Correlation of revealed preference (decision thresholds) with stated preference (average video ratings) for Experiments 1 and 2. (I, K, M) Individual subject correlation coefficients between revealed and stated preferences in Experiment 1. (J, L, N) Individual subject correlation coefficients between revealed and stated preferences in Experiment 2.

### 3.4 Predicting Behavioural Reliability

We calculated z-scored standard deviations of the task variables across sessions as an index of reliability of the task over time. We then used a series of LASSO models to identify any questionnaire predictors of behavioural reliability. In addition, a PCA of our questionnaire data (see Supplementary Materials) indicated two underlying components – internalising symptoms (PC1) and externalising symptoms (PC2). We ran a series of linear models to identify which PCs were significant predictors of behavioural reliability (see Table 6).

**Table 6.**
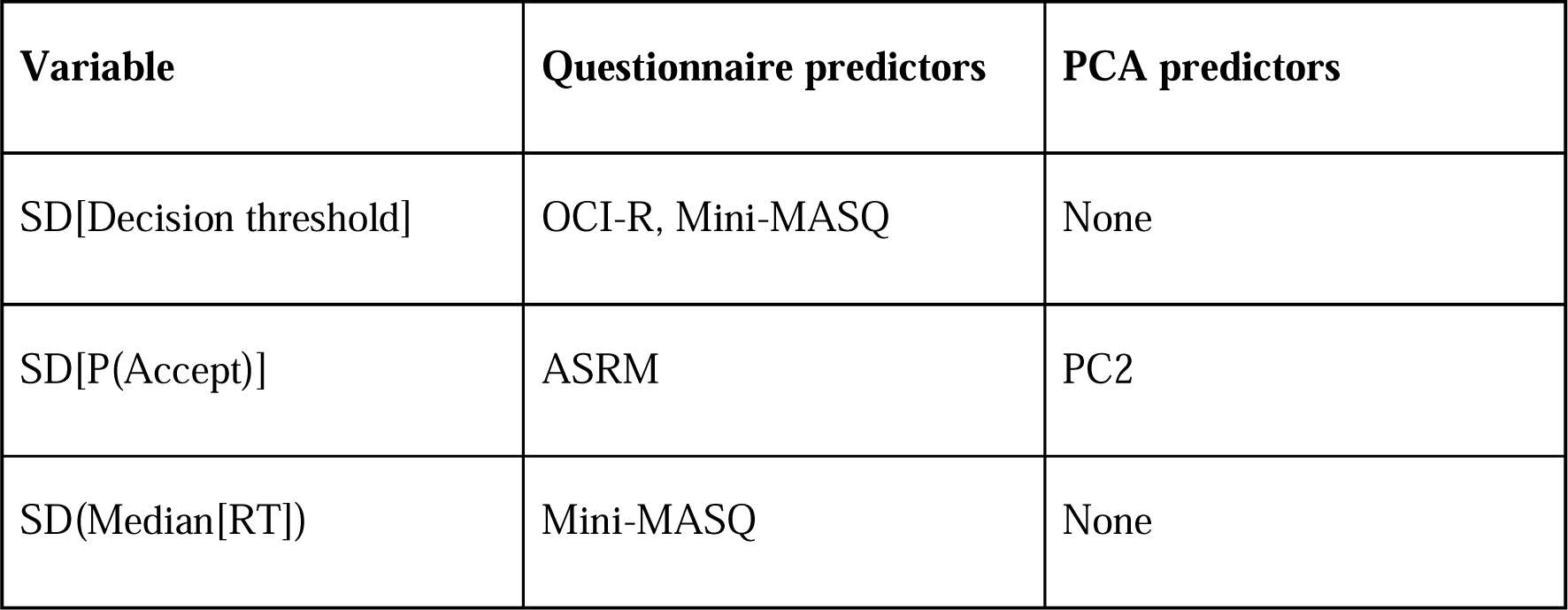
Predictors of behavioural stability across sessions in Experiment One.

| Variable | Questionnaire predictors | PCA predictors |
| --- | --- | --- |
| SD[Decision threshold] | OCI-R, Mini-MASQ | None |
| SD[P(Accept)] | ASRM | PC2 |
| SD(Median[RT]) | Mini-MASQ | None |

LASSO regression of the standard deviation of decision thresholds across sessions identified obsessive-compulsive (OCI-R), *F*_(1, 37)_ = 6.83, *p* = .013, *R*^2^ = .05, and mood/anxiety (Mini-MASQ), *F*_(1, 37)_ = 7.52, *p* = .009, *R*^2^ = .05, symptoms as predictors. For the LASSO model of P(Accept) variability, mania (ASRM) was a predictor, *F*_(1, 36)_ = 5.33, *p* = .028, *R*^2^ = .09. In addition, PC2 was a significant predictor of P(Accept) variability, *F*_(1, 36)_ = 5.19, *p* = .030, *R*^2^ = .226. Finally, mood and anxiety symptoms (Mini-MASQ) were identified as a significant predictor of variability in decision times across sessions, *F*_(1, 38)_ = 6.15, *p* = .018, *R*^2^ = .21. Predictors of behavioural reliability in Experiment One are plotted in Figures 5A-E and 5G. The predictive utility of mania symptoms (ASRM), *F*_(1, 36)_ = 5.87, *p* = .020, *R*^2^ = .12, and PC2 scores, *F*_(1, 36)_ = 5.04, *p* = .030, *R*^2^ = .11, on P(Accept) variability was replicated in Experiment Two (see Figures 4F and 4H).

**Figure 4.**
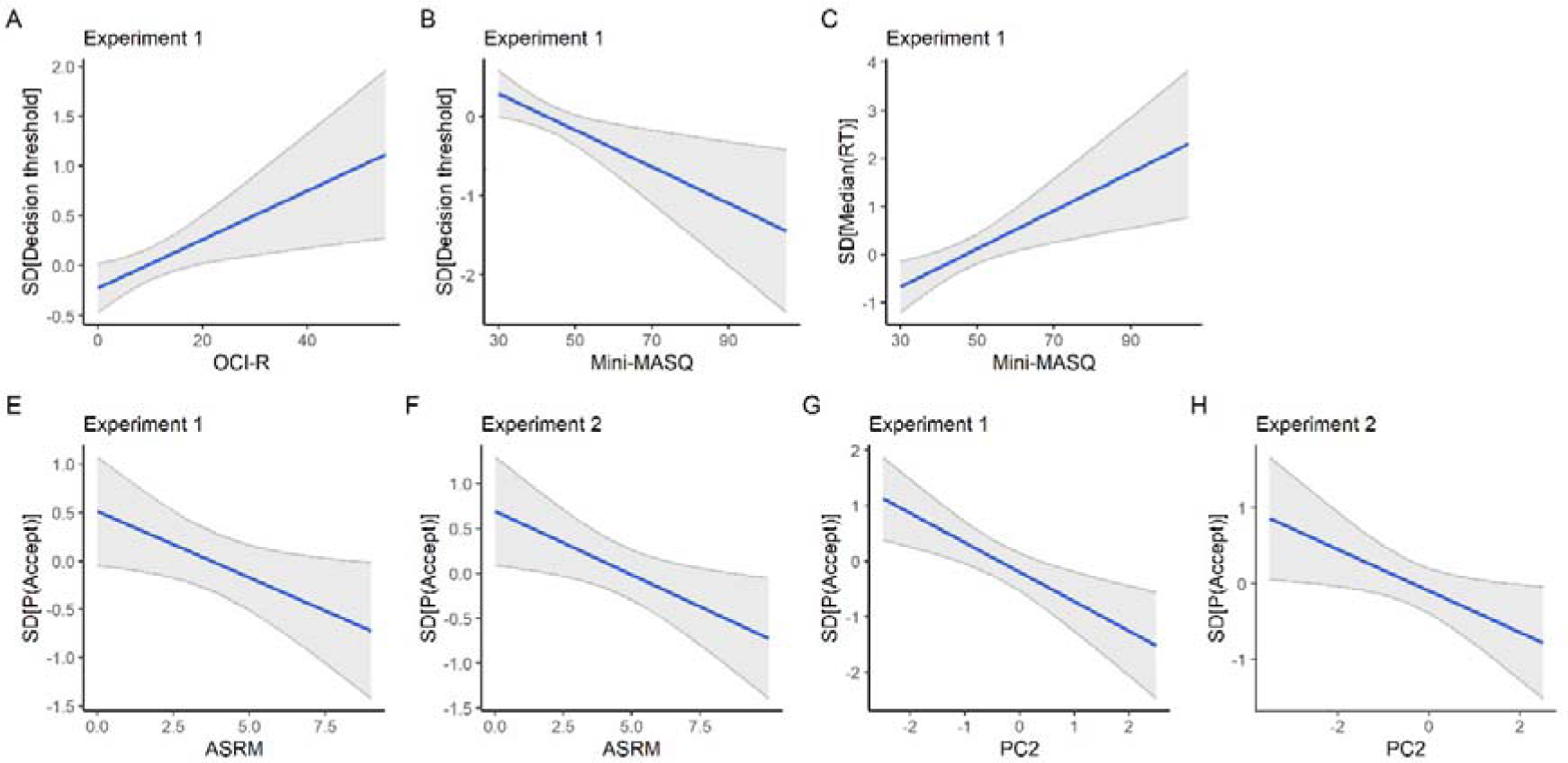
Model predictions of behavioural stability across sessions in Experiment One (A-E) and Experiment Two (F-G).

**Figure 5.**
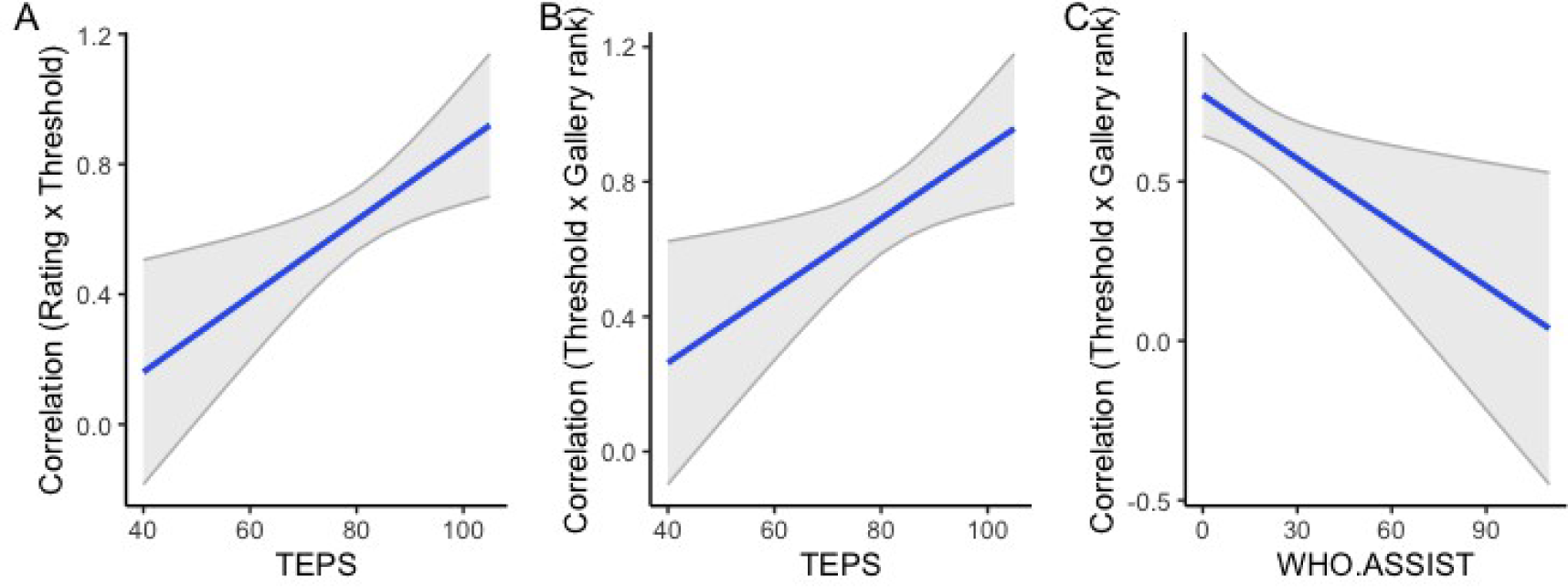
Model predictions of hedonic reliability in Experiment One.

### 3.5 Predicting Hedonic Reliability

For each participant, we calculated correlation coefficients between each combination of their decision thresholds, average video ratings, and explicit gallery rankings. With these correlation coefficients, we ran a series of LASSO models to identify which questionnaires could predict hedonic reliability. We also ran a series of linear models to identify which PCs were significant predictors of these correlation coefficients (see Table 7).

**Table 7.** Predictors of hedonic reliability in Experiment One.

| Correlation variables | Questionnaire predictors | PCA predictors |
| --- | --- | --- |
| Decision threshold x average video rating | TEPS | None |
| Average video rating x Explicit gallery ranking | None | None |
| Decision threshold x Explicit gallery ranking | TEPS, WHO-ASSIST | None |

For the correlations of average video ratings with decision thresholds, pleasure-seeking (TEPS) was identified as a predictor, *F*_(1, 111)_ = 7.38, *p* = .007, *R*^2^ = .07. In addition, for the correlations of decision thresholds with explicit video rankings, pleasure-seeking (TEPS), *F*_(1, 109)_ = 4.83, *p* = .030, *R*^2^ = .06, and drug use (WHO.ASSIST), *F*_(1, 109)_ = 6.42, *p* = .013, *R*^2^ = .06, were retained as predictors. Predictors of hedonic reliability in Experiment One are plotted in Figure 5. We ran the same linear models examining hedonic reliability with data from our validation sample. None of the effects observed in Experiment One were replicated in Experiment Two.

### 3.6 Behavioural Predictions

Using LASSO regression, no questionnaires were retained as predictors of average decision threshold, P(Accept), average video ratings, or median decision time. Similarly, using linear mixed models, PC1, PC2, and PC3 were not significant predictors of average decision threshold, P(Accept), or median decision latency. However, PC2 was a significant predictor of average video ratings, *F*_(1, 107)_ = 14.46, *p* < .001, *R*^2^ = .129, and that prediction replicated in Experiment Two, *F*_(1, 109)_ = 7.26, *p* = .008, *R^2^* = .06. The model predictions across Experiments One and Two are shown in Figure 6.

**Figure 6.**
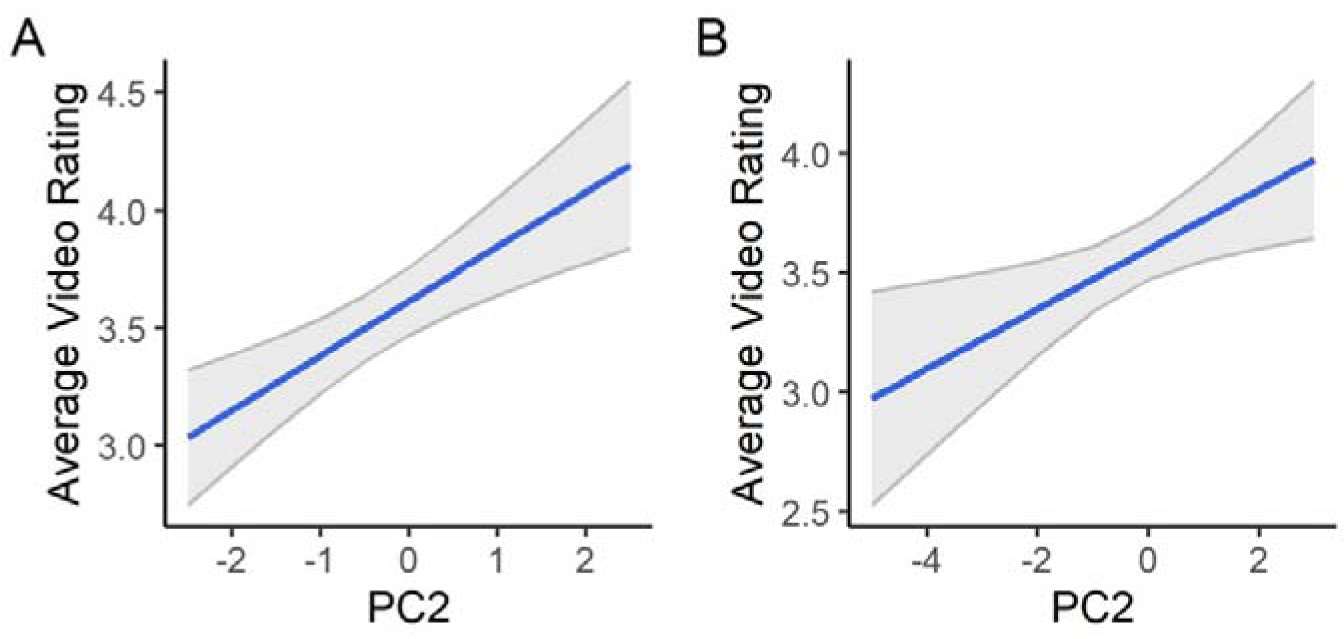
Model predictions of behaviour in Experiment One (A) and Experiment Two (B).

## 4.0 Discussion

### 4.1 Foraging as an etiological approach to investigate psychopathology

Quantifying underlying neurocognitive profiles may serve as a useful approach to understand the etiological commonalities to psychopathology and improve therapies to address them. A key challenge to consider in adopting this approach is that there is a lack of demonstrably reliable and predictive behavioural assays that can index clinically relevant neurocognitive profiles. Psychophysical tasks are often applied to measure specific facets of cognition at the expense of capturing multiple overlapping constructs. As an attempt to address this, complex behaviour is increasingly being studied in psychiatric research to understand typical and pathological cognition as it relates to naturalistic behaviour (Hasler, 2012; Kishida et al., 2010). In particular, foraging paradigms can provide a computational account of naturalistic behaviour, allowing an examination of maladaptive decisional processes that may support psychopathology (Adams et al., 2016; Huys et al., 2016; Redish & Gordon, 2016).

The WebSurf foraging paradigm offers a mechanistically relevant, translational platform to investigate decision making, which is particularly applicable to psychiatric research because translational accounts of neurocognitive processing are critical for developing successful interventions. Therefore, an obvious next step is examining whether the WebSurf paradigm has utility in objectively quantifying psychopathology. Behavioural patterns within the WebSurf task have previously been linked to psychopathology. For example, Abram et al. (2019) applied a risk component to the WebSurf task, where the time delay imposed before reward was presented as a range of delays and the true delay was only known by the participant once they had accepted an offer. In non-risk trials, participants were informed of the exact time delay before reward when choosing to accept or reject an offer.

Individuals who were higher in externalising disorder vulnerability, and thus more vulnerable to addiction, were more likely to accept a risky offer when they had received a bad outcome on the previous trial and valued bad outcomes after risky decisions as more pleasurable. Two separate variants of the WebSurf task have been applied in humans – one in which participants navigate a virtual maze for video rewards (Movie Row) and one in which participants receive candy as rewards, rather than videos (Candy Row). In the Movie Row task, participants with a BMI > 25 were more likely to accept offers for which the delay was longer than their decision threshold (Huynh et al., 2021). Therefore, there may be some relation of task behaviour with psychobiological processes. However, it is unclear whether task behaviour can be predictive of more broad psychopathological profiles. Furthermore, while task behaviour remains relatively stable over multiple sessions of Restaurant Row, WebSurf retest reliability has not been demonstrated.

### 4.2 The challenge of remote behavioural assessment

The first challenge in validating the neurocognitive utility of the WebSurf task is administering it online in uncontrolled and unsupervised environments. Similar behavioural patterns have been observed in the WebSurf task across samples administered remotely and in-person (Huynh et al., 2021; Kazinka et al., 2021; Redish et al., 2022). However, participants may engage in task-irrelevant distractions to countermand the boredom elicited by the imposed time delays for reward. To ensure data validity, our platform monitored the status of the browser window in which the task was running and could detect if the window was minimised, or browser tabs had been switched (see Supplementary Materials). If an inactive task window was detected, the task quit, and participants were excluded from further participation. A large proportion of participants failed this attention check, despite being informed that they would be unable to complete the task if they did not keep their task window active. Across our administrations of the task, the failure rate was as high as 35.7%. Furthermore, while we employed additional post-hoc attention checks, we cannot confirm that participants were not engaging in other task-irrelevant distractions. In addition, there was attrition in participants returning to complete repeated administrations of the task within the allocated timeframe and the completion rate of all study components was as low as 40%.

Increasing that engagement and retention would be an important consideration for studies seeking to deploy WebSurf or similar attentional tasks in clinical populations at scale.

Interestingly, depression, mania, and schizotypy symptoms could predict study completion in Experiment One and when accounting for depression and mania symptoms, the predictive value of schizotypy on study completion was validated in Experiment Two.

### 4.3 Stability of task behaviour and the exploration/exploitation trade-off

WebSurf behaviour is generally stable over time. Our Bayesian analysis for the most part indicated strong- to very-strong evidence against an effect of session number on behavioural parameters. An exception to this is the number of threshold violations which showed some evidence of systematic change across repeated sessions. This suggests that the optimality of choices made in the task may be less stable over time. Our calculation of ICCs over repeated sessions further support the reliability of task behaviour. When considering all three sessions of the task, ICCs of task parameters showed good reliability, with the exception of P(Accept) and the number of threshold violations. However, further examination of individual session combinations shows where the stability of parameters breaks down. Across all behavioural parameters, and across both our exploratory and validation samples, ICCs showed excellent retest reliability when considering only the second and third administrations of the task.

Behavioural stability was consistently lower when considering ICCs across the first and second, and first and third administrations of the task. This suggests that the first session produces behaviour which is prone to change. At initial exposure to the task, participants may spend more time in an exploratory approach where they explore unfamiliar options for reward (Addicott et al., 2017). This exploration of options and rewards likely involves an internal formation of preference, willingness to wait, and strategy for reward maximisation, which is reflected in task behaviour. During this time, participants may form rules for decisions which can heuristically optimise the decision-making process but may also manifest as overly-rigid and habitual behaviour common to psychopathology. The stability of behaviour at future exposure to the task suggests that participants largely switch to an exploitation strategy for reward maximisation. A future point of consideration may address whether momentary task behaviour can indicate state-like changes in exploration and exploitation. Given the noradrenergic system’s role in the exploration/exploitation trade-off, pupillometry could be a particularly useful in capturing these state-like changes as it is thought to index locus coeruleus activity (Joshi et al., 2016), which plays a key role in the release of noradrenaline throughout the central nervous system (Foote et al., 1983; Jepma & Nieuwenhuis, 2011).

Furthermore, our finding that initial task performance is less reliable but remains stable in future task administrations is informative for future applications of the WebSurf task.

These data suggest that in obtaining stable behavioural performance in WebSurf, it may be useful for participants to complete an initial exposure to the task prior to completing the entire assay, because subsequent measures are likely to be more stable and retention was much greater between the second and third sessions. The stability of task behaviour in repeated administrations supports its utility as a clinical tool, but there is a large proportion of attrition which poses a threat to its application. In addition, we observed a predictive utility of mood and anxiety symptoms and externalising symptoms on P(Accept) variability. These changes in behavioural stability suggest there may be underlying differences in how individuals with psychiatric symptoms approach the task, which may similarly reflect shifts in exploratory versus exploitative strategies for reward maximisation.

### 4.4 Hedonic reliability

Correlating stated preferences with revealed preferences can provide an indication of hedonic reliability. When unreliable, this may suggest that conscious deliberative processes are disrupted or that there is an alteration in affective motivation (Barch et al., 2017). Across our samples, the consistency between stated and revealed preferences was generally high, although there was variation between individuals. None of the administered questionnaires could predict hedonic reliability in a replicable manner. This suggests that hedonic reliability is a construct that is not reliably captured by traditionally psychometric scales.

### 4.5 The link from symptoms to behaviour

In our exploratory analysis, we observed a relationship between externalising symptoms and task behaviour, reflected in P(Accept). This is consistent with the alterations of behaviour in the risk-taking variant of the WebSurf task for those with high trait externalising symptoms (Abram, Redish, et al., 2019). However, our metric of externalising symptoms was not directly taken from an externalising symptom measurement, but was derived via our questionnaire battery PCA. Therefore, these results suggest there is some predictive utility of task behaviour on broad psychopathological symptom patterns. While the effect replicated in our confirmatory sample, there was a large number of tested parameters in our exploratory sample which did not show any predictive utility. These findings suggest that the behavioural parameters we investigated here may capture dimensions of psychological variance that are different than that of traditional psychometric rating scales. While we note that method variation may attribute to the modest correlation of extracted parameters with self-reported distress, a critical point is that task behaviour may reflect psychopathological variance that is not captured by self-report. Importantly, many emerging therapies for psychiatric disorders directly target aberrant decisional processing (Amidfar et al., 2019; Basu et al., 2023; Fisher et al., 2010; Haber et al., 2020; Subramaniam et al., 2014). Behavioural metrics that can repeatably quantify complex cognitive constructs are thus necessary to develop these technologies further. Given that WebSurf is demonstrably reliable, assesses multiple aspects of decision-making simultaneously, and is clinically scalable, it offers a promising paradigm to assess and track psychiatric phenomena.

### 4.6 Conclusion

In summary, we have validated a mechanistically relevant, reverse translatable, and clinically viable assessment of multiple decision-making systems. Psychopathology is increasingly thought to arise from dysfunctional information processing which is often highly variable between individuals, but many neurocognitive assessments suppress the variance between individuals. This limits their ability to capture the heterogeneity prevalent in psychiatric phenomena. The WebSurf task addresses this by integrating complex, naturalistic behaviour to assess mechanistically relevant multi-system function and track behavioural variance between individuals. Here we demonstrate that behaviour within individuals is highly stable after initial exposure to the task, supporting its utility as a tool to track clinical phenomena.

Behavioural parameters extracted from the task can predict anhedonic or externalising symptomology, but importantly, task parameters may measure dimensions of psychological variance that are not captured by traditional rating scales. An integration of these etiological paradigms to quantify psychiatric phenomena is thus relevant to understand and treat psychiatric disorders by providing a more holistic platform to measure the complex systems underpinning them.

## Acknowledgements

We thank A. David Redish for critical intellectual discussions around the overall design and concept of the project, as well as for contributions to the original WebSurf task. We acknowledge the technical support provided by the University of Minnesota Department of Psychiatry & Behavioral Sciences Computerized Psychiatric Assessment Suite (COMPAS) development team, and particularly Dr. Karrie Fitzpatrick.

## Author Contributions

A.N.M., C.R.P.S., A.M.III., and A.S.W. conceived and designed research; A.N.M. performed experiments; C.R.P.S provided regulatory support; A.N.M. analysed data; A.N.M., A.M.III., and A.S.W. interpreted results of experiments; A.N.M. prepared figures; A.N.M. drafted manuscript; A.N.M., A.M.III., and A.S.W. edited and revised manuscript; A.N.M., C.R.P.S., A.M.III., and A.S.W. approved the final version of manuscript.

## 6.0 Supplementary Materials

### 6.1 Websurf Behavioural Screening

A variant of the WebSurf task presents an attention check during the wait period, which requires participants to click an on-screen button when it changes colour (Kazinka et al. 2021). While this element may be beneficial in encouraging participant engagement and therefore experience of the delay especially during un-proctored online administration of the task, this attention check may itself serve as a distraction from the wait period, which in turn changes behaviour. For instance, although humans performing WebSurf without an attention check show a sunk costs fallacy (Redish et al., 2022; Sweis et al., 2018), adding the attention check eliminates this effect (Kazinka et al. 2021). Therefore, since the inclusion of attention checks can modify behaviour, we chose not to include this attention check in our task. Rather, our task would quit if it was detected that a participant had minimized their browser window or changed browser tabs. If this was detected, participants were marked as having failed the task, and were excluded from the study.

We also screened the collected behavioural data for evidence that subjects were inattentive during their participation. We flagged session data for rejection if their median decision latency was > 3000 ms, because it has been demonstrated that most decisions in the task are made within three seconds (Abram et al., 2016). Thus, a median reaction time longer than this would suggest that participants were not sufficiently engaged in the task. We also flagged session data for rejection if participants chose to accept all offers, or if they chose to reject all offers. Participants were excluded from analysis if all their completed sessions indicated poor engagement in the task via these behavioural checks. In addition, subjects may show an “inverted” threshold for which, contrary to expected behaviour, they tend to stay for offers above their decision threshold and skip offers below their decision threshold (Huynh et al., 2021; Kazinka et al., 2021). This may be indicative that subjects were using a task-irrelevant distraction during the wait period and as such, subjects were flagged for rejection if the inverted model fit their data better than the expected model.

### 6.2 Behavioural Parameters

We calculated several parameters of behaviour in the WebSurf task. These are decision thresholds, P(Accept), average video ratings, median decision times, and number of threshold violations across video gallery ranks and across sessions for the sample of participants who completed all three sessions of WebSurf. These variables (collapsed across video galleries) are plotted across sessions in Supplementary Figure S1 shows the parameters as a function of session number, aggregated across video galleries. Supplementary Figure S2 shows these parameters as a function of session number and explicit video gallery ranking.

**Supplementary Figure S1.**
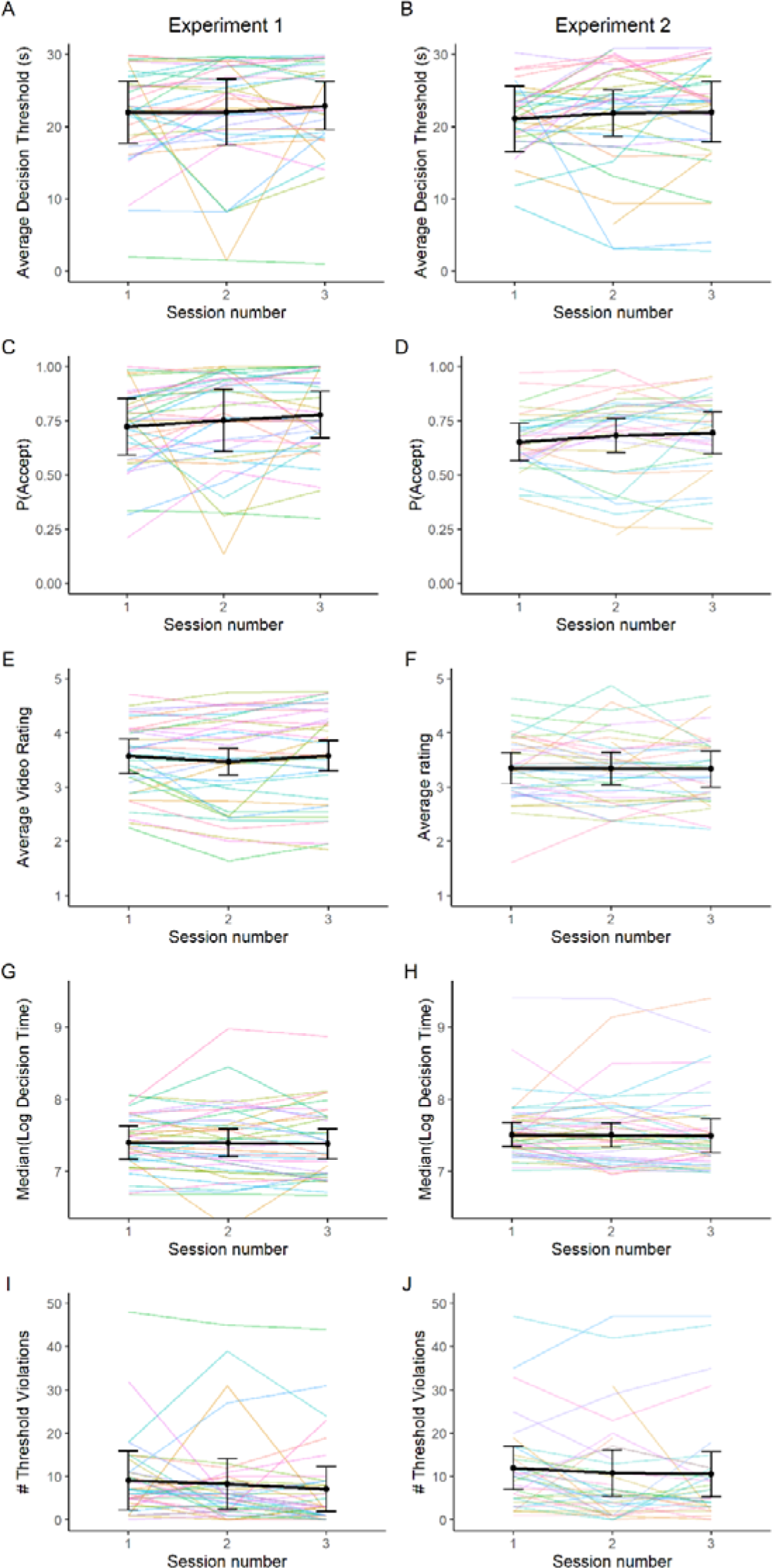
Behavioural parameters across three sessions of the WebSurf task in Experiment One (left panels) and Experiment Two (right panels) and collapsed across video galleries. Coloured lines represent individual subject measurements. Points represent the mean across subjects for each session number. Error bars represent standard deviation.

**Supplementary Figure S2.**
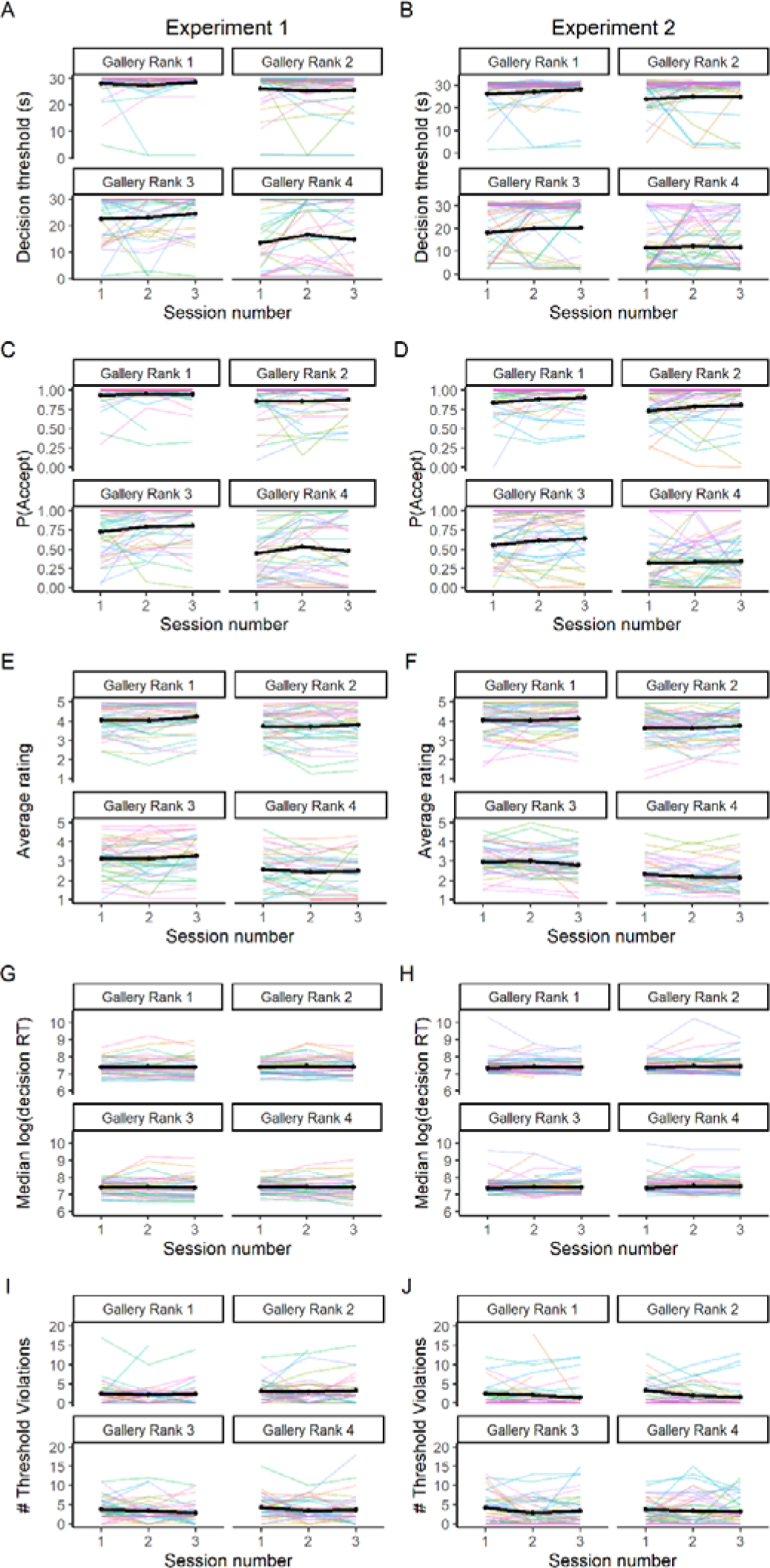
Behavioural parameters across three sessions of the WebSurf task in Experiment One (left panels) and Experiment Two (right panels) and across video galleries according to explicit gallery ranking. Coloured lines represent individual subject measurements. Points represent the mean across subjects for each session number. Error bars represent standard deviation.

### 6.3 Intraclass Correlations

We calculated ICCs of these variables as a metric of behavioural reliability across repeated administrations of the task. We examined all session combinations (i.e., reliability across sessions one and two, across sessions one and three, across sessions two and three, and across all sessions). Supplementary Figure S3 shows ICCs for task behavioural parameters and across video galleries for Experiment One, and Supplementary Figure S4 shows these data for Experiment Two.

**Supplementary Figure S3.**
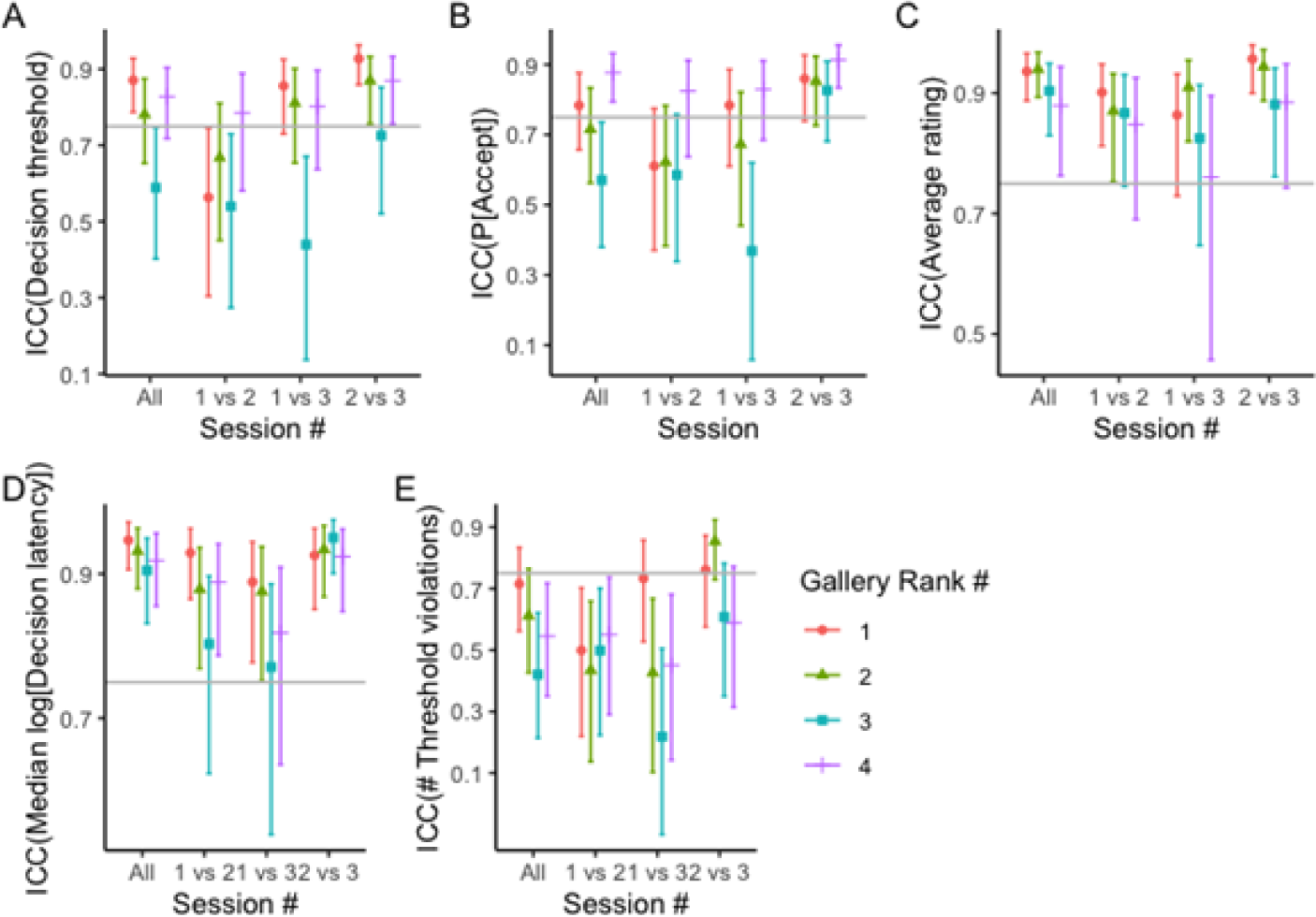
Intraclass correlations of behavioural parameters between sessions and across gallery rankings in Experiment One.

**Supplementary Figure S4.**
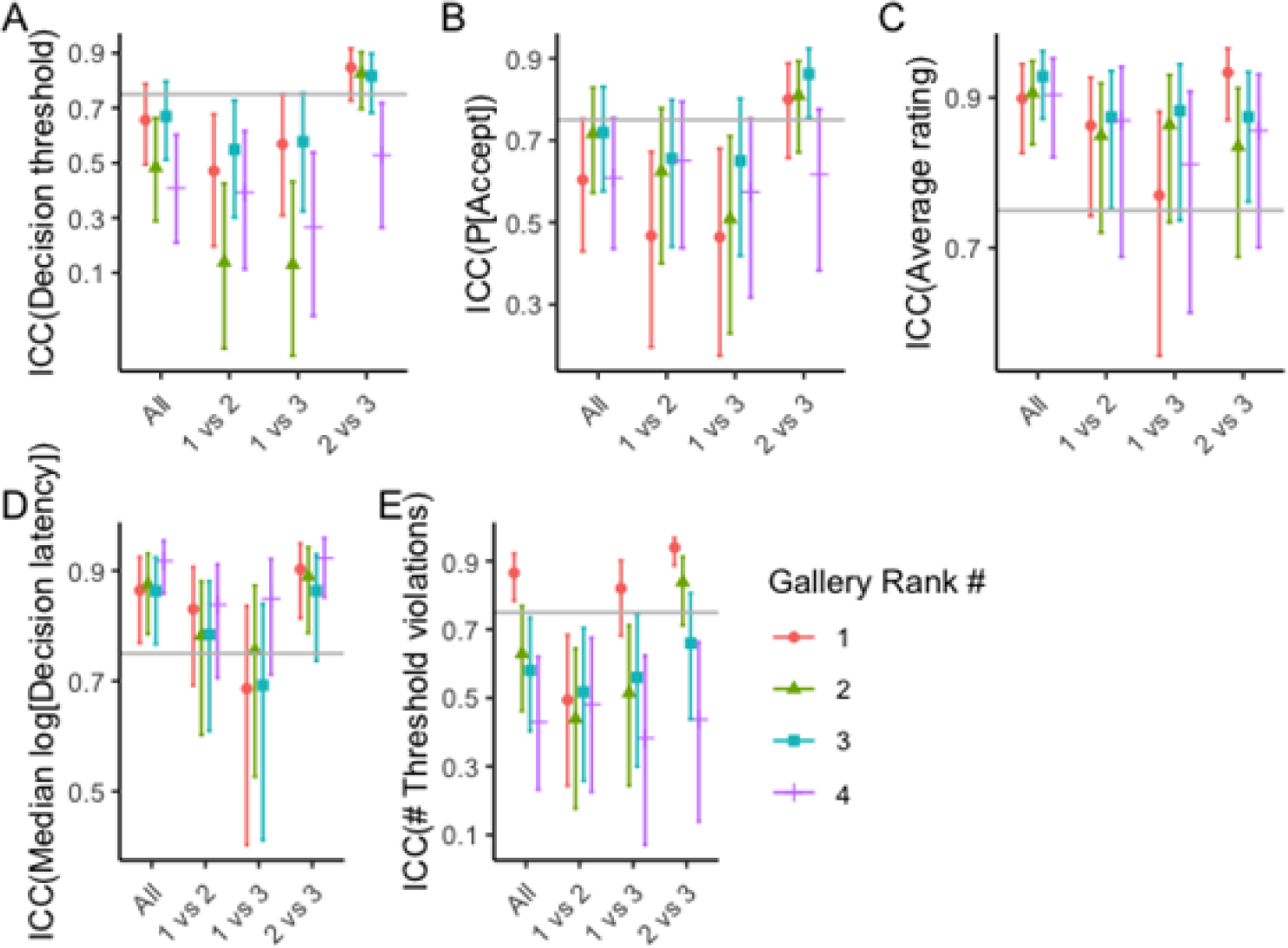
Intraclass correlations of behavioural parameters between sessions and across gallery rankings in Experiment Two.

### 6.4 Psychometric Questionnaires

At recruitment into the study, participants completed a battery of self-report rating scales. The questionnaire battery included the Altman Self-Rating Mania Scale (ASRM; Altman et al., 1997), the Alcohol Use Disorders Identification Test (AUDIT; Saunders et al., 1993), the Aberrant Salience Inventory (ASI; Cicero et al., 2010), the Centre for Epidemiological Studies – Depression (CES-D; Radloff, 2016), the Daily Sessions, Frequency, Age of Onset, and Frequency of Cannabis Use (DFAQ-CU; Cuttler & Spradlin, 2017), the Eating Atttitudes Test (EAT-26; Garner et al., 1982), the Mini Mood and Anxiety Symptoms Questionnaire (Mini-MASQ; Watson et al., 1995), the Obsessive Compulsive Inventory – Revised (OCI-R; Foa et al., 2002), the Snaith-Hamilton Pleasure Scale (SHAPS; Snaith et al., 1995), the Barratt Impulsivity Scale (BIS-11; Patton et al., 1995), the Schizotypal Personality Questionnaire – Brief (SPQ-B; Raine & Benishay, 1995), the Temporal Experience of Pleasure Scale (TEPS; Gard et al., 2006), and the World Health Organization Alcohol, Smoking, and Substance Involvement Screening Test (WHO-ASSIST; WHO ASSIST Working Group, 2002). Descriptions of each measure and their subscales are found in Supplementary Table S1.

**Supplementary Table S1.** Questionnaires contained in battery.

| **Name** | **Core Reference** | **Measured Construct** | **Number of items** | **Subscales** |
| --- | --- | --- | --- | --- |
| Altman Self-Rating Mania Scale (ASRM) | Altman et al. (1997) | Severity of manic and hypomanic symptoms | 5 | ☐ Positive mood<br>☐ Self-confidence<br>☐ Sleep patterns<br>☐ Speech patterns<br>☐ Motor activity |
| Alcohol Use Disorders Identification Test (AUDIT) | Saunders et al. (1993) | Alcohol use | 10 | ☐ Alcohol consumption<br>☐ Drinking behaviors<br>☐ Alcohol-related problems |
| Aberrant Salience Inventory (ASI) | Cicero et al. (2010) | Unusual or incorrect valuation of stimuli as being salient or significant – related to psychosis and psychotic disorders | 29 | ☐ Increased significance<br>☐ Senses sharpening<br>☐ Impending understanding<br>☐ Heightened emotionality<br>☐ Heightened cognition |
| Centre for Epidemiological Studies – Depression (CES-D) | Radloff (2016) | Severity of symptoms associated with depression | 20 |  |
| Daily Sessions, Frequency, Age of Onset, and Frequency of | Cuttler and Spradlin (2017) | Cannabis use | 24 | ☐ Frequency<br>☐ Age of onset of cannabis use |
| Cannabis Use (DFAQ-CU) |  |  |  | ? Quantity |
| Eating Attitudes Test (EAT-26) | Garner et al. (1982) | Symptoms characteristic of eating disorders | 26 | ? Dieting<br>? Bulimia and food preoccupation<br>? Oral control |
| Mini Mood and Anxiety Symptoms Questionnaire (Mini-MASQ) | Watson et al. (1995) | Severity of symptoms related to depression and anxiety | 26 | ? General distress<br>? Anxiety symptoms<br>? Depression symptoms |
| Obsessive Compulsive Inventory – Revised (OCI-R) | Foa et al. (2002) | Severity of symptoms related to obsessive-compulsive disorder | 18 | ? Washing<br>? Obsessing<br>? Hoarding<br>? Ordering<br>? Checking<br>? Neutralizing |
| Snaith-Hamilton Pleasure Scale (SHAPS) | Snaith et al. (1995) | Experience of pleasure vs. anhedonia | 14 | ? Interest/pastimes<br>? Social interaction<br>? Sensory experience<br>? Food/drink |
| Barratt Impulsivity Scale (BIS-11) | Patton et al. (1995) | Impulsivity | 30 | Six first-order factors<br>? Attention<br>? Motor impulsiveness<br>? Self-control<br>? Cognitive complexity<br>? Perseverance<br>? Cognitive instability |
|  |  |  |  | Three second-order factors<br>☐ Attentional impulsivity<br>☐ Motor impulsivity<br>☐ Non-planning impulsivity |
| Schizotypal Personality Questionnaire – Brief (SPQ-B) | Raine and Benishay (1995) | Severity of symptoms related to schizotypal personality disorder | 22 | ☐ Cognitive-perceptual<br>☐ Interpersonal<br>☐ Disorganized |
| Temporal Experience of Pleasure Scale (TEPS) | Gard et al. (2006) | Experience of pleasure vs. anhedonia | 18 | ☐ Anticipatory pleasure<br>☐ Consummatory pleasure |
| World Health Organization Alcohol, Smoking, and Substance Involvement Screening Test (WHO-ASSIST) | WHO-ASSIST Working Group (2002) | Usage of alcohol and other drugs | 120 | ☐ Tobacco<br>☐ Alcohol<br>☐ Cannabis<br>☐ Cocaine<br>☐ Stimulants<br>☐ Inhalants<br>☐ Sedatives<br>☐ Hallucinogens<br>☐ Opioids<br>☐ Other drugs |

### 6.5 Principal Component Analysis

Prior to running PCA on questionnaire data, one participant was excluded from Experiment One because they failed more than one attention check item in the questionnaire battery. PCA of the questionnaire data from Experiment One revealed four PCs with eigenvalues > 1.

However, the fourth component explained less than 10% of variance in questionnaire scores. Therefore, we retained only the first three components for further analyses. Supplementary Figure S5A-B shows correlations and histograms of questionnaire total scores in Experiment One. In Supplementary Figure S5E, we present the percentage of variance accounted for by each of the PCs in Experiment One. The first PC predominantly contained contributions from questionnaires relating to internalizing symptoms and psychopathology, such as depression (CES-D, Mini-MASQ), anxiety (Mini-MASQ), and schizotypy (SPQ.B). The second PC contained contributions from externalizing symptoms related to pleasure seeking (TEPS, SHAPS) and mania (ASRM). The third PC received strongest contributions from questionnaires related to alcohol (WHO.ASSIST, AUDIT) and other drug (WHO.ASSIST, DFAQ.CU) use.

Supplementary Figure S5C-D shows correlations and histograms of questionnaire total scores in Experiment Two and Supplementary Figure S5F, shows the percentage of variance accounted for by each of the PCs in Experiment Two. The structure of the PCs in Experiment Two were similar to those in Experiment One. PC1, reflecting internalizing symptoms, received the strongest contributions from the same questionnaires as Experiment One. PC2, representing externalizing symptoms, received strongest contributions from symptoms related to pleasure-seeking (TEPS/SHAPS), but unlike Experiment One, also received strong contributions from symptoms relating to psychosis (ASI). PC3, representing alcohol and other drug use, received the strongest contributions from questionnaires related to alcohol and other drug use (WHO.ASSIST, DFAQ.CU), but also from pleasure-seeking (TEPS).

Given the overall consistency between the PCs extracted across our exploratory and validation samples, and to enable replication of the effects we observed in Experiment One, we applied the same PCA rotations from our Experiment One exploratory sample to the questionnaire scores obtained from our Experiment Two validation sample. This produced individual subject PC scores which were comparable across the two samples. Histograms of individual subject PC scores across the exploratory and validation samples are shown in Supplementary Figure S6. In addition, the rotations applied by the PCAs of questionnaire scores in Experiment One and Experiment Two are shown in Supplementary Figures S7 and S8.

**Supplementary Figure S5.**
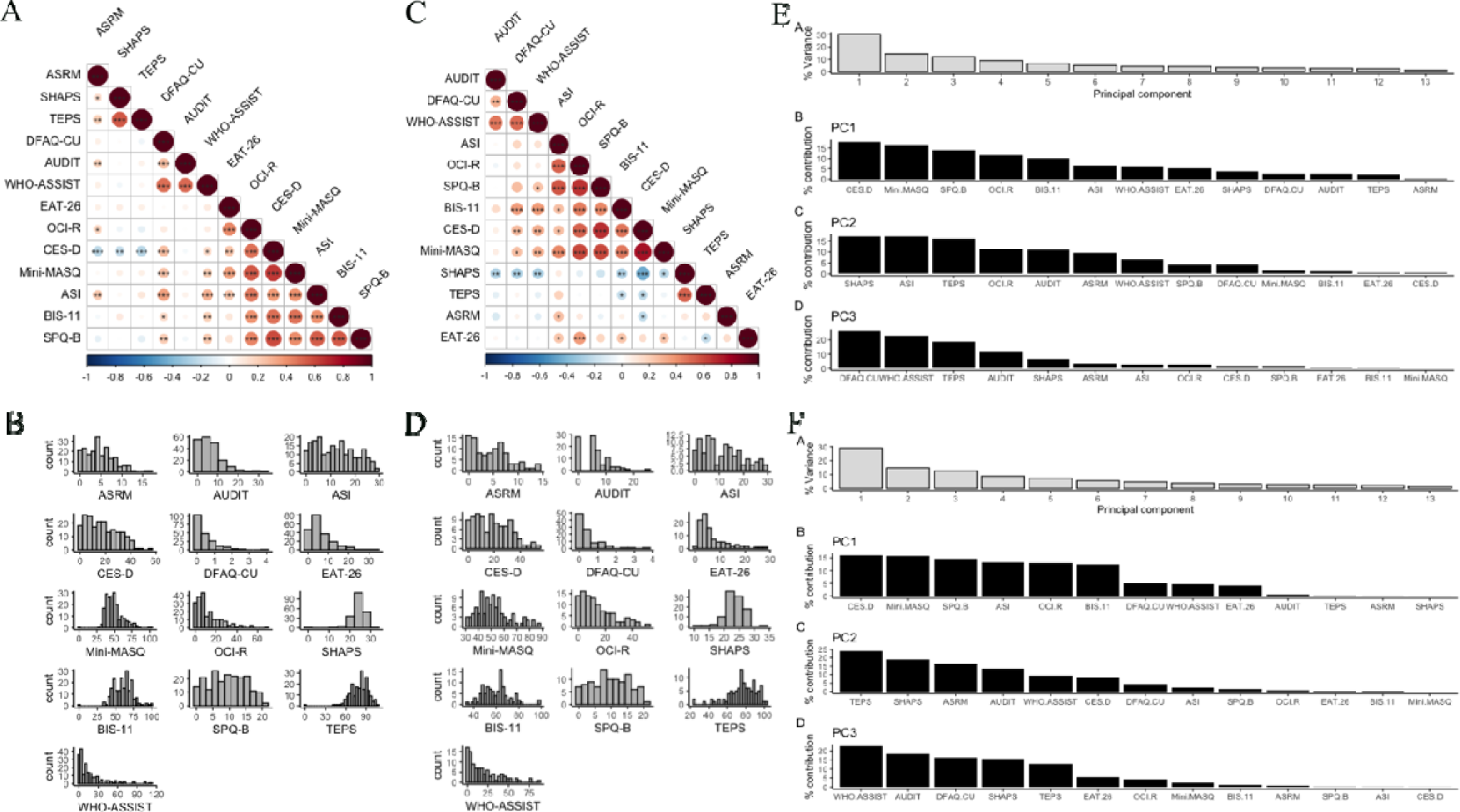
(A) Correlations of questionnaire total scores in Experiment One. (B) Histograms of questionnaire total scores in Experiment One. (C) Correlations of questionnaire total scores in Experiment Two. (D) Histograms of questionnaire total scores in Experiment Two. (E) Percentage of questionnaire variance accounted for by each of the Principal Components in Experiment One. (F) Percentage of questionnaire variance accounted for by each of the Principal Components in Experiment Two.

**Supplementary Figure S6.**
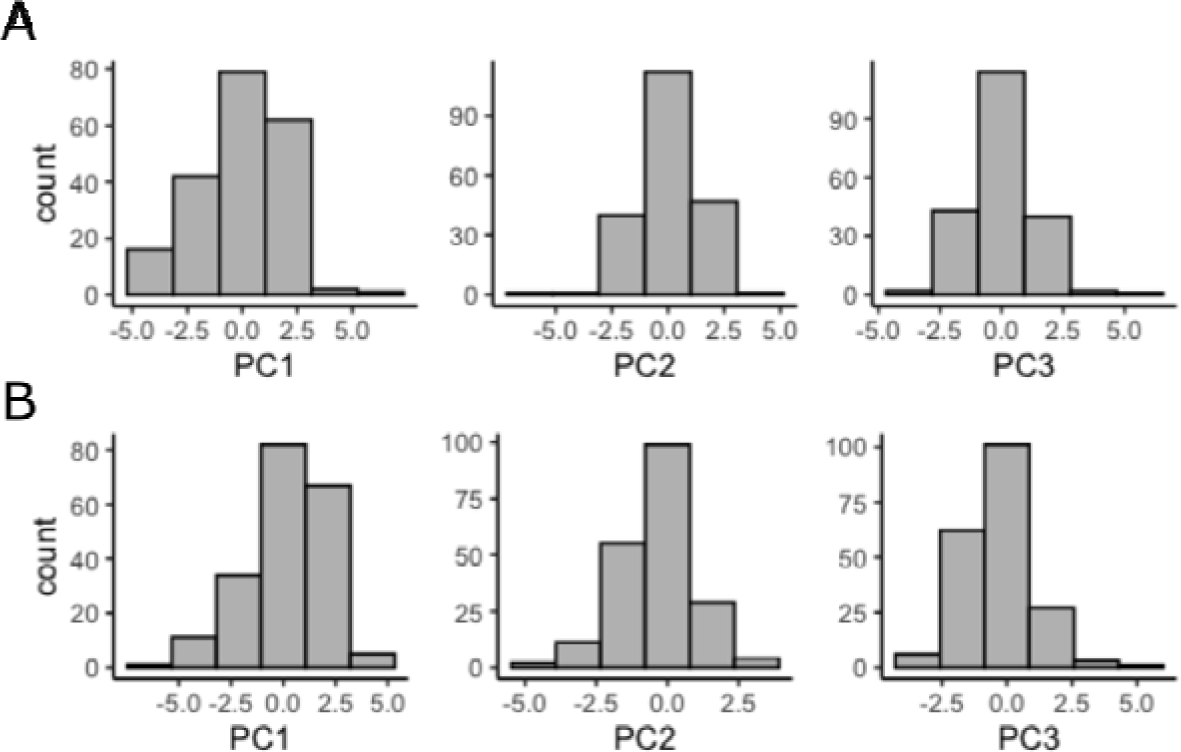
Individual subject Principal Component scores in Experiment One (A) and Experiment Two (B).

**Supplementary Figure S7.**
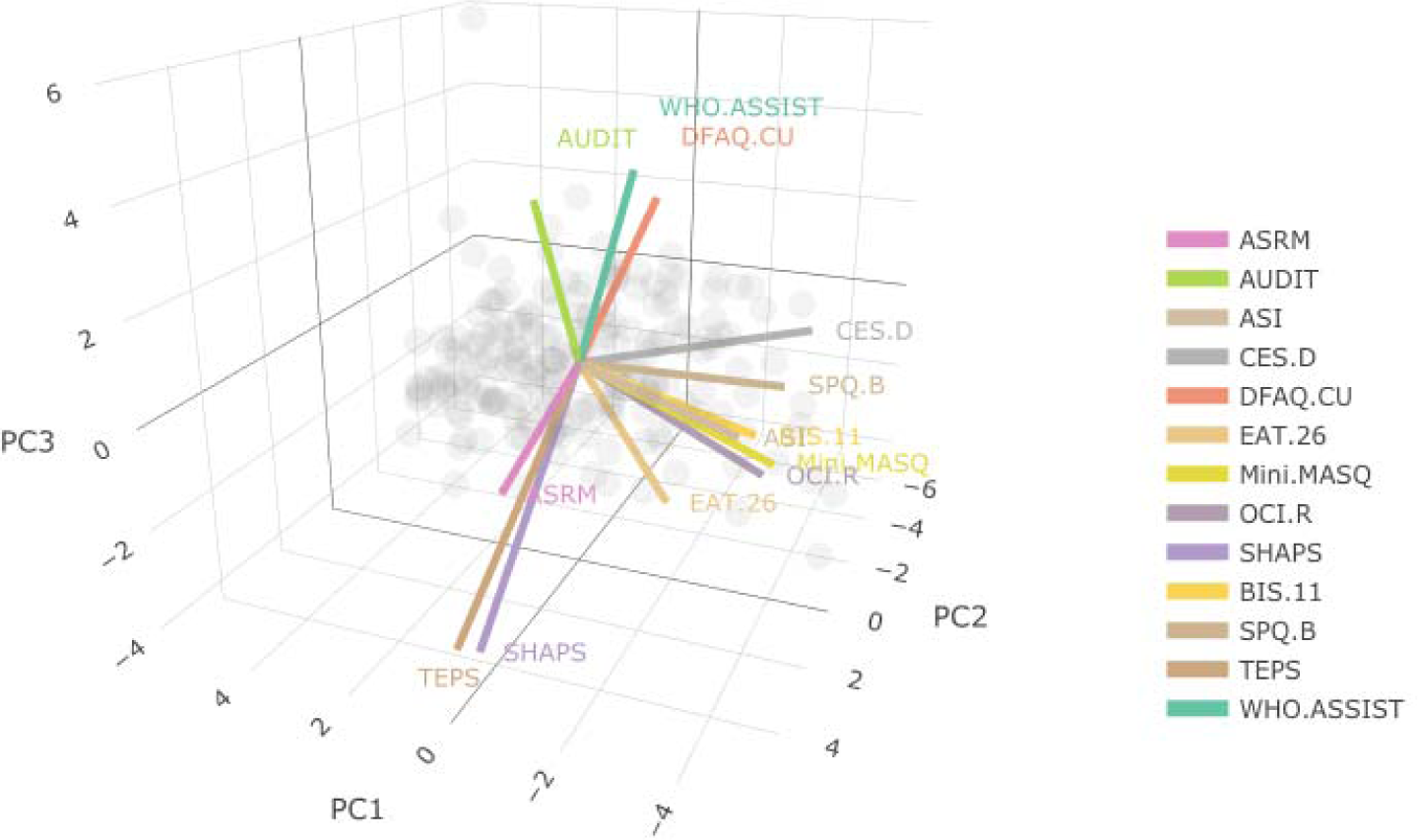
Rotations applied by Principal Component Analysis of questionnaire scores in Experiment One.

**Supplementary Figure S8.**
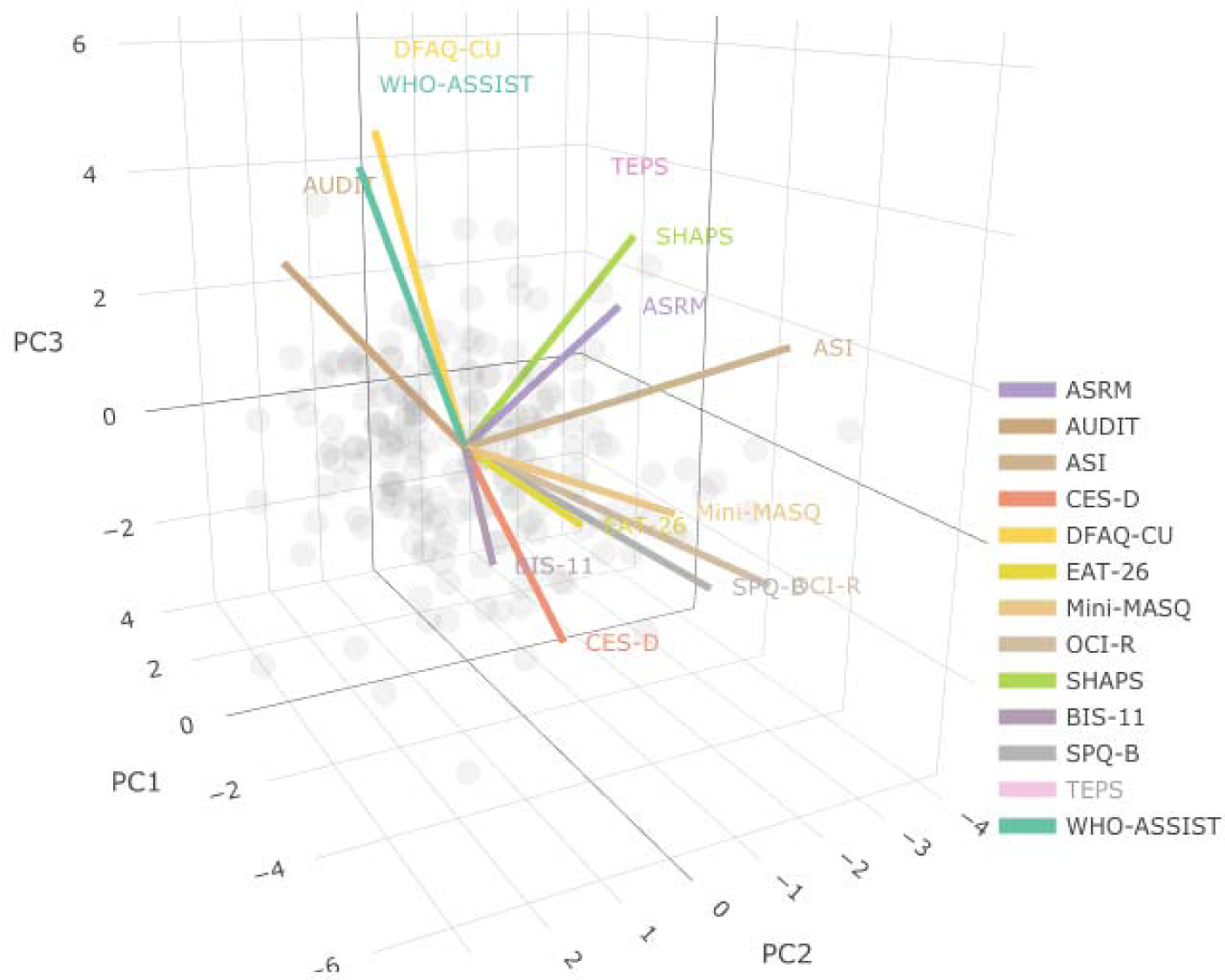
Rotations applied by Principal Component Analysis of questionnaire scores in Experiment Two.

### 6.6 Predicting Study Completion

We calculated a binary variable representing whether participants completed (1) or did not complete (0) all components of the study. To assess the predictive utility of task behaviour on individual symptom profiles, we selected predictive questionnaire scores using LASSO regression. As a supplement, we examined whether task behaviour better predicts broader psychopathological profiles by entering PCs into logistic regressions. A LASSO logistic regression with questionnaire scores as predictors identified mania (ASRM), depression (CES-D), and schizotypal personality (SPQ) as predictors of study completion. The logistic regression identified a negative association of both mania (ASRM; OR[95% CI] = 0.82[0.71, 0.95]) and depression (CES-D; OR[95% CI] = 0.94[0.89, 0.99]) with probability of study completion, whereas the direction was positive for schizotypy (SPQ; OR[95% CI] = 1.17[1.05, 1.31]).

The predictive value of mania (ASRM) and depression (CES-D) symptoms on study completion failed to replicate in Experiment Two. However, schizotypy (SPQ) was again predictive of study completion with a positive direction, OR[95% CI] = 1.22[1.04, 1.44]. Of note, in both Experiments, schizotypy symptoms were only predictive of study completion when accounting for mania (ASRM) and depression (CES-D) symptoms in the model. As an exploratory analysis, we conducted the same logistic regression with only schizotypy (SPQ) symptoms as a predictor, which was not significant. Model predictions across the two experiments are plotted in Supplementary Figure S9.

**Supplementary Figure S9.**
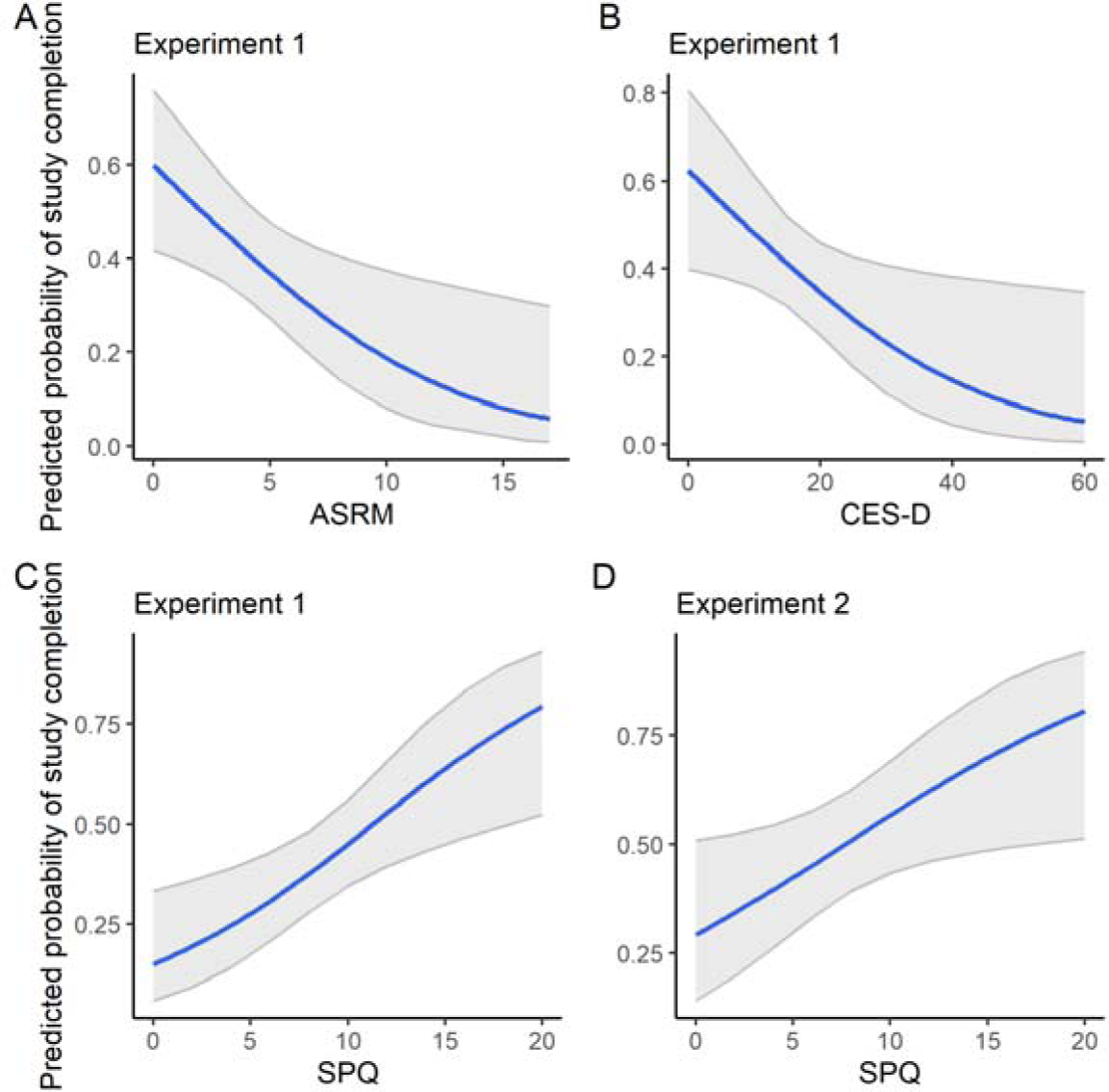
Psychiatric symptom predictors of study completion in Experiment One (A-C) and Experiment Two (D).

